# Transcriptional Readthrough at *Atf4* Locus Suppresses *Rps19bp1* and Impairs Heart Development

**DOI:** 10.1101/2025.08.26.672495

**Authors:** Zengming Zhang, Tongbin Wu, Zeyu Chen, Danni Chen, Zhengyu Liang, Christopher Adams, Yu Gu, Mao Ye, Fhujjen Barroga, Sylvia Evans, Xiaohai Zhou, Ju Chen

## Abstract

**BACKGROUND:** Activating Transcription Factor 4 (ATF4) functions as a transcriptional regulator in various cell types and tissues under both physiological and pathological conditions. While previous studies have linked ATF4 activation with promoting cardiomyocyte (CM) death in dilated cardiomyopathy (DCM), atrial fibrillation, and heart failure, its role in developing CMs remains unexplored.

**METHODS:** We generated multiple distinct CM-specific (*Atf4^cKO(e2/3/pA)^* and *Atf4^cKO(e2)^*) and global *Atf4* knockout (*Atf4^7del/7del^* and *Atf4^1ins/1ins^*) mouse models targeting different *Atf4* regions, as well as cardiomyocyte-specific deletion of *Rps19bp1* to study cardiac phenotypes. Detailed morphological and molecular analyses were performed.

**RESULTS:** *Atf4^cKO(^*^e2/3*/pA)*^ (targeting exon 2-3 including the polyadenylation signal (polyA)) mice exhibited severe cardiac defects and died before E17.5, likely due to ectopic activation of p53 signaling pathway resulting from *Rps19bp1* downregulation, a potent suppressor of p53. Further investigation revealed that deleting the polyA signal of *Atf4* in *Atf4^cKO(e2/3/pA)^* mice led to transcriptional readthrough, resulting in the formation of an *Atf4*-*Cacna1i* fusion transcript and *Rps19bp1* downregulation. To avoid readthrough while abolishing ATF4 function, we introduced small indels into exon 3 of *Atf4* in mice (*Atf4^7del/7del^* and *Atf4^1ins/1ins^*), which showed normal *Rps19bp1* expression and cardiac morphology. Importantly, CM-specific deletion of *Rps19bp1* recapitulated the cardiac defects and transcriptional change seen in *Atf4^cKO(e^*^2^*^/3/pA)^* mice.

**CONCLUSIONS:** We found that the downregulation of *Rps19bp1*, not loss of ATF4 function, underlying the cardiac phenotypes in *Atf4^cKO(e2/3/pA)^* mice. The reduced expression of *Rps19bp1* in *Atf4^cKO(e2/3/pA)^* mice is likely due to the unintentional deletion of *Atf4* polyA signal and subsequent transcriptional readthrough, underscoring the essential role of RPS19BP1, not ATF4, in cardiac development. Consistent *Rps19bp1* downregulation has been observed in other tissue-specific *Atf4* knockout models utilizing the *Atf4^fl(e2/3/pA)^* allele, suggesting that previously reported *Atf4* KO phenotypes may result from *Atf4* transcriptional readthrough effects. These findings reveal a locus-dependent transcriptional interference mechanism and emphasize the importance of avoiding confounding cis effects in genetically engineered models.

**TRANSLATIONAL PERSPECTIVE:** Our findings clarify ATF4’s role in heart development by showing that cardiac defects in cardiomyocyte-specific ATF4 knockout mice—using a widely employed floxed ATF4 line—result from unintended downregulation of RPS19BP1 caused by transcriptional readthrough. This shifts the focus from ATF4 to RPS19BP1, a key regulator of p53 activity, as a potential driver of cardiac developmental abnormalities. Clinically, these insights caution against misinterpretation of genetic knockout models and highlight RPS19BP1 as a promising target for congenital heart disease and related cardiac dysfunctions, with potential implications for future therapies.

## Introduction

Activating Transcription Factor 4 (ATF4), also known as cAMP-Response Element Binding Protein 2 (CREB2), is a basic region leucine zipper transcription factor involved in physiological responses to various stressors including hypoxia, endoplasmic reticulum stress, amino acid deprivation, oxidation, and mitochondrial stress ^1–4^. ATF4 has been implicated in the pathogenesis of diverse diseases, including liver steatosis^5^, insulin resistance^6^, Alzheimer’s disease^7^, tumor formation^8^ and skeletal muscle atrophy^9–12^. In the cardiovascular system, ATF4 overexpression-induced ER stress contributes to vascular atherosclerotic calcification and CM cell death^13, 14^. Moreover, ATF4 upregulation is observed in patients with dilated cardiomyopathy (DCM), atrial fibrillation, and heart failure^15–18^, suggesting its role as a disease-associated TF in adult hearts. Furthermore, ATF4 plays a critical role in development of various tissues. Global *Atf4*-deficient mice exhibit perinatal lethality, accompanied by impaired hematopoiesis^19, 20^, subfertility^21^, defective lens formation^22, 23^, abnormal osteoblast differentiation and bone homeostasis^24–27^, and erythroid differentiation defects, leading to hypoplastic anemia^28^. Despite these findings, our understanding of ATF4 functions in embryonic development, especially heart development, remains incomplete.

To investigate the role of ATF4 in heart development, we generated cardiomyocyte (CM)-specific *Atf4* knockout (KO) mice (*Atf4^cKO(e2/3/pA)^*) by crossing the widely used *Atf4* floxed (*Atf4^fl(e2/3/pA)/fl(e2/3/pA)^*) mice, which target exon 2 and exon 3 including the polyA sequence. *Atf4^cKO(e2/3/pA)^*mice exhibited thinner ventricular walls and died perinatally with abnormal activation of p53 pathway. Further studies revealed that p53 was increased at the protein level, likely due to the downregulation of *Rps19bp1*, a potent suppressor of p53^29^. Intriguingly, *Rps19bp1* and *Cacna1i*, two of the most dysregulated genes in *Atf4^cKO(e2/3/pA)^*mice, are located adjacent to *Atf4* in the genome, implying that their dysregulation may result from genetic disruption of the *Atf4* locus, rather than the loss of ATF4 function. RNA-seq data further revealed that exon 1 of *Atf4* and exon 2 of *Cacna1i* were aberrantly spliced together, generating an *Atf4*-*Cacna1i* fusion transcript. In addition, IGV visualization of RNA-seq coverage showed widespread RNA read coverage across the intergenic region between *Atf4* and *Cacna1i*, which indicate the presence of *Atf4* transcriptional readthrough in *Atf4^cKO(e2/3/pA)^* samples.

Transcriptional readthrough refers to the phenomenon in which the transcriptional machinery fails to terminate properly at polyadenylation or other transcriptional stop signals, resulting in continued transcription into downstream regions^30–32^. *Rps19bp1* is located between *Atf4* and *Cacna1i* while transcribed from the opposite strand, *Atf4* transcriptional readthrough likely suppressed *Rps19bp1* expression due to collision of transcriptional machineries^33, 34^. To minimize potential artifacts caused by transcriptional readthrough, we generated a novel CM-specific *Atf4* KO mouse line (*Atf4^cKO(e2)^*) targeting exon 2 of *Atf4*. Unexpectedly, *Atf4^cKO(e2)^* mice produced a truncated ATF4 protein. To avoid the possibility that the truncated ATF4 protein retains residual activity, we generated two *Atf4* mutant lines (*Atf4^7del/7del^* and *Atf4^1ins/1ins)^*) carrying frameshift mutations in exon 3 that abolished ATF4 protein expression. Although most mutants died perinatally, both lines exhibited normal cardiac morphology and transcriptome profiles with normal expression of *Rps19bp1* and *Cacna1i*. Collectively, these findings demonstrate that ATF4 is not required for embryonic heart development and adult heart function.

To investigate whether the phenotypes observed in *Atf4^cKO(e2/3/pA)^* were due to downregulation of *Rps19bp1* caused by transcriptional readthrough, we generated *Rps19bp1* CM-specific KO mice (*Rps19bp1^cKO^*) and observed that *Rps19bp1^cKO^* exhibited identical cardiac defects, reduced survival, and highly correlated gene expression alterations as seen in *Atf4^cKO(e2/3/pA)^* mice, providing strong evidence that the observed phenotypes resulted from *Atf4* readthrough-caused *Rps19bp1* downregulation, rather than the loss of ATF4 function.

Taken together, our findings suggest that *Atf4* transcriptional readthrough, rather than loss of ATF4 function, caused the cardiac development defects in *Atf4^cKO(e2/3/pA)^* mice. Importantly, *Rps19bp1* downregulation has also been noted in other tissues-specific KO or global-knockout mice. Our results suggest that phenotypic and molecular changes observed in these studies may similarly stem from *Rps19bp1* downregulation consequent to *Atf4* transcriptional readthrough. Our study underscores the importance of re-evaluating previous findings on *Atf4* and emphasizes the need for caution in future research to avoid potential misinterpretations of results

## Methods

### Animals

All animal procedures were performed in accordance with the National Institutes of Health Guide for the Care and Use of Laboratory Animals and were approved by the Institutional Animal Care and Use Committee of the University of California, San Diego. *Atf4^fl(e2/3/pA)/fl(e2/3/pA)^*mice were previously generated by inserting two LoxP sites flanking exons 2 and 3 ^12^. *Atf4^fl(e2)/fl(e2)^* mice were generated by inserting two LoxP sites flanking exons 2. *Atf4^7del/7del^* and *Atf4^1ins/1ins^* models were generated by introducing a small 7bp deletion and a 1bp insertion in the exon3 of *Atf4* gene, respectively. All mice are of C57BL/6N background. *Rps19bp1^fl/fl^*mice(C57BL/6JGpt) were purchased in GemPharmatech company (Strain ID: T022391). To obtain heart tissue for further molecular and cellular analysis, pregnant mice are euthanized by ketamine injection (100mg/kg) (Dechra) followed by cervical dislocation. Genotypes of mice were confirmed by polymerase chain reaction (PCR) analysis using embryonic yolk sac or tail extracts and primer sequences for genotypes were shown in Table S1.

### Echocardiography

Mouse echocardiography was performed as described ^35^. Briefly, the animals were initially anesthetized with 5% isoflurane (VETone, 502017) for one minute, followed by maintenance at 1% throughout the examination. The anterior chest wall was shaved and then Nair was applied to remove any remaining hair. Small needle electrodes were inserted into one upper and one lower limb for simultaneous electrocardiogram. Transthoracic echocardiography (M-mode and 2-dimensional) was conducted using the VisualSonics, FUJIFILM, Vevo 2100 ultrasound system with a linear transducer 32-55MHz. Various cardiac parameters including heart rate (HR), left ventricular end-diastolic dimensions (LVED) and left ventricular end-systolic dimensions (LVESD), end-diastolic interventricular septal thickness (IVSd) and LV posterior wall thickness (LVPWd) were assessed from the LV M-mode tracing. Percentage fractional shortening (%FS) served as an indicator of systolic cardiac function.

### Western Blot

Mouse hearts tissues were harvested and snap-frozen in liquid nitrogen. Total protein extracts were prepared by homogenization of hearts in RIPA buffer (Thermo Fisher Scientific, 89901). Protein concentration was determined using a Micro BCA Protein Assay Kit (Thermo Fisher Scientific). Tissue lysate was mixed with 4x LDS sample buffer and 10x Reducing Agent (Life Technologies) and incubated for 10 minutes at 70°C. Protein lysates were separated on Bolt 4% to 12% SDS-PAGE gels (Thermo Fisher, NW04125BOX) and transferred onto polyvinylidene fluoride (PVDF) membranes (Bio-Rad, 1620177) overnight at 4 °C. After blocking in blocking buffer (TBS containing 0.1% Tween-20 and 5% dry milk) for 1h, the membranes were incubated overnight at 4 °C with primary antibodies: p53 (leica biosystem, NCL-L-p53-CM5p), GAPDH (Santa Cruz, sc-32233). Membranes were then washed with TBST and incubated with HRP-conjugated secondary anti-rabbit IgG (Dako – Agilent, P0448/ P044801-2) or anti-mouse IgG (Dako – Agilent, P0447/ P044701-2) for 2 hours at room temperature. Immunoreactive protein bands were visualized using enhanced chemiluminescence (ECL) reagent (Bio-Rad) and captured by Bio-Rad ChemiDoc Imaging System.

### Histology and Immunofluorescence

Embryonic mouse hearts were dissected at various developmental stages and fixed in 4% PFA overnight at 4°C. Fixed hearts were then dehydrated in 5%, 10%, 15%, 20% sucrose and embedded Tissue-Tek in OCT (Sakura, 4583). Tissues were sectioned at 8 μm sections using a Leica CM 3050S cryostat (Leica Microsystems). For histology, the mouse sections were then stained with Hematoxylin and Eosin (H&E) using a standard protocol (Procedure No. HT110, Sigma). Images were captured using a Hamamatsu NanoZoomer 2.0HT Slide Scanning System. For immunofluorescence, the mouse heart sections were blocked with antibody block buffer (5% BSA, 0.2% Tween-20 in PBS) for one hour and then incubated with primary antibodies NKX2-5 (Santa Cruz Biotech, SC8697) or Ki67 (Abcam, ab15580) or Caspase 3 (CST, 9661S) overnight at 4°C in a humidified chamber. After washed three times with PBST (PBS with 0.1% Triton X-100), the sections were incubated with secondary antibodies for 2 hours at room temperature, and then sections were counterstained with DAPI and mounted with ProLong™ Gold Antifade Mountant medium (Invitrogen, P36930). Images were captured using an Olympus FluoView FV1000 Confocal Microscope.

### TUNEL Assay

Terminal deoxynucleotidyl transferase-mediated dUTP nick end labeling (TUNEL) assays were executed on heart sections with an in situ Cell Death Detection Kit (Roche, 11684795910) that labeled the nuclei of dying cells with green fluorescence. Cardiomyocyte nuclei were co-stained with an Nkx2.5 (NK2 homeobox 5) antibody (Santa Cruz Biotech, SC8697).

### EdU Labelling

To assess cell proliferation in embryonic hearts, pregnant female mice were injected intraperitoneally with EdU (5-ethynyl-20-deoxyuridine, Invitrogen) two hours before embryo dissection. Following cryosectioning of embryonic hearts, EdU-positive cells were detected using the Click-iT assay (Invitrogen) with Alexa Fluor 647 azide according to the manufacturer’s instructions. This was followed by primary and secondary antibody incubations per standard immunofluorescence procedures. Images were captured using an Olympus FluoView FV1000 Confocal Microscope.

### RNA Sequencing

Total RNA was extracted from E11.5 mouse ventricular hearts using TRIzol (Invitrogen) following the manufacturer’s instructions. The quantity and quality of purified RNA was assessed by Agilent 2100 Bioanalyzer. RNA integrity numbers (RIN) were in the range of 6.1 to 10. RNA-Seq libraries (n = 4 mice per genotype) were prepared using an Illumina TruSeq stranded mRNA kit according to manufacturer’s instructions. High-output 150-cycle paired-end sequencing was performed with an Illumina HiSeq 6000 sequencer at the UCSD Institute for Genomic Medicine (IGM) core facility to a sequencing depth of 30-70 million reads per sample. RNA-Seq analyses were conducted in Linux (AlmaLinux release 8.9), R (version 4.2.1) and Python (version 3.6.8). Trim Galore (https://www.bioinformatics.babraham.ac.uk/projects/trim_galore/) was used for quality and adapter trimming as well as quality control. The mouse reference genome sequence and gene annotation data, mm10, were downloaded from UCSC Genome Browser (https://hgdownload.cse.ucsc.edu/goldenpath/mm10/). The RNA-Seq reads were mapped onto the genome using Hisat2 (version 2.2.1). SAMtools (version 1.3.1) was employed to sort the alignments, and the FeatureCounts python package was employed to count reads per gene. The DESeq2 R Bioconductor package (54) was used to normalize read counts and identify DEGs, using FDR-adjusted P values (Benjamini-Hochberg method) < 0.05 and fold change (FC) >1.5 as thresholds. Gene Ontology (GO) Enrichment analysis of DEGs was performed using DAVID (https://david.ncifcrf.gov/), using a cutoff of p value < 0.01. GSEA was performed using clusterProfiler R package and software GSEA.

### Data availability

All data supporting the findings of this study are provided in the main text or the supplementary materials. The bulk RNA-seq data generated in this study have been deposited in NCBI’s Gene Expression Omnibus (GEO) under the accession numbers GSE303780, GSE303775 and GSE303929.

### Quantitative Real-Time Polymerase Chain Reaction

Total RNA was isolated from frozen embryonic hearts using TRIzol solution (Invitrogen), Approximately one microgram RNA was used for reverse transcription with M-MLV Reverse Transcriptase according to the manufacturer’s instructions (Promega, M1701). Quantitative real-time polymerase chain reaction reactions were performed using iTaq Universal SYBR Green Supermix (Bio-Rad, 1725120) on CFX Opus 96 Real-Time Polymerase Chain Reaction System (Bio-Rad). The relative expression levels of target genes were determined using the comparative CT method (ΔΔCt method) and normalized to the mRNA levels of the housekeeping gene *Polr2a*. Primer sequences for qRT-PCR were shown in Table S1.

### Statistical Analysis

ImageJ software was used for image analysis. All statistical analyses were performed using GraphPad Prism 8 software. Student’s t-tests were employed to compare means between two groups. Data are presented as mean ± SEM. The significance of differences between groups was determined as indicated in each figure legend. P-values less than 0.05 were considered significant and denoted as follows: *p < 0.05, **p < 0.01, ***p < 0.001, ****p < 0.0001.

## Results

### Cardiomyocyte-specific deletion of *Atf4* impairs embryonic heart development

To investigate the role of ATF4 in the heart, we first generated global *Atf4* knockout (*Atf4^gKO(e2/3/pA)^*) mice by crossing *Sox2-Cre* transgenic mice with a widely used *Atf4* floxed line (targeting exons 2–3, including the polyA signal) (*Atf4^fl(e2/3/pA)/fl(e2/3^*^/*pA*)^) ^12^, which resulted in embryonic lethality before embryonic day 8.5 (E8.5) (Figure S1), suggesting that ATF4 is required for early embryo survival. To study the cell autonomous function of ATF4 in cardiomyocytes (CMs), we created CM-specific *Atf4* knockout (KO) mice (*Atf4^fl(e2/3/pA)/fl(e2/3/pA)^*; *Xmlc2-Cre^+/-^*, hereafter *Atf4^cKO(e2/3/pA)^*) by crossing *Atf4^fl(e2/3/pA))/fl(e2/3/pA)^* mice with *Xenopus laevis myosin light-chain 2 (Xmlc2)-Cre* mice, which specifically drives Cre expression in CMs as early as E7.5 ^36–39^. *Atf4^cKO(e2/3^*^/*pA*)^ mice displayed abnormal heart morphology at E12.5 (Figure 1A-B), and most *Atf4^cKO(e2/3/pA)^*embryos died between E15.5 and E17.5 (Figure S2A). Hematoxylin and eosin (H&E) staining revealed that *Atf4^cKO(e2/3/pA)^* mice have thinner compact myocardium in both ventricles from E12.5 onward (Figure 1C-F). To determine whether the reduced thickness was due to decreased CM proliferation or increased CM apoptosis, we used specific cell cycle markers (Ki-67 for cell cycle activity and EdU for DNA synthesis) to assess the percentage of proliferating CMs, and cleaved caspase 3 (cCSP3) immunofluorescence and TUNEL assays to evaluate CM apoptosis in *Atf4^cKO(e2/3/pA)^* mice and littermate controls at E11.5, a stage preceding the appearance of abnormal heart morphology in *Atf4^cKO(e2/3/pA)^* hearts (Figure 1G). EdU was administered 2 hours prior to embryos collection to label cells undergoing DNA synthesis. We observed a significant reduction in the proliferation rate of CMs in *Atf4^cKO(e2/3/pA)^*mice (Figure 1G-H), while there was no difference in CM apoptosis (Figure S2B-C), indicating that the thinner LV and RV compact myocardium in *Atf4^cKO^* hearts was primarily due to CM proliferation defects.

**Figure 1.**
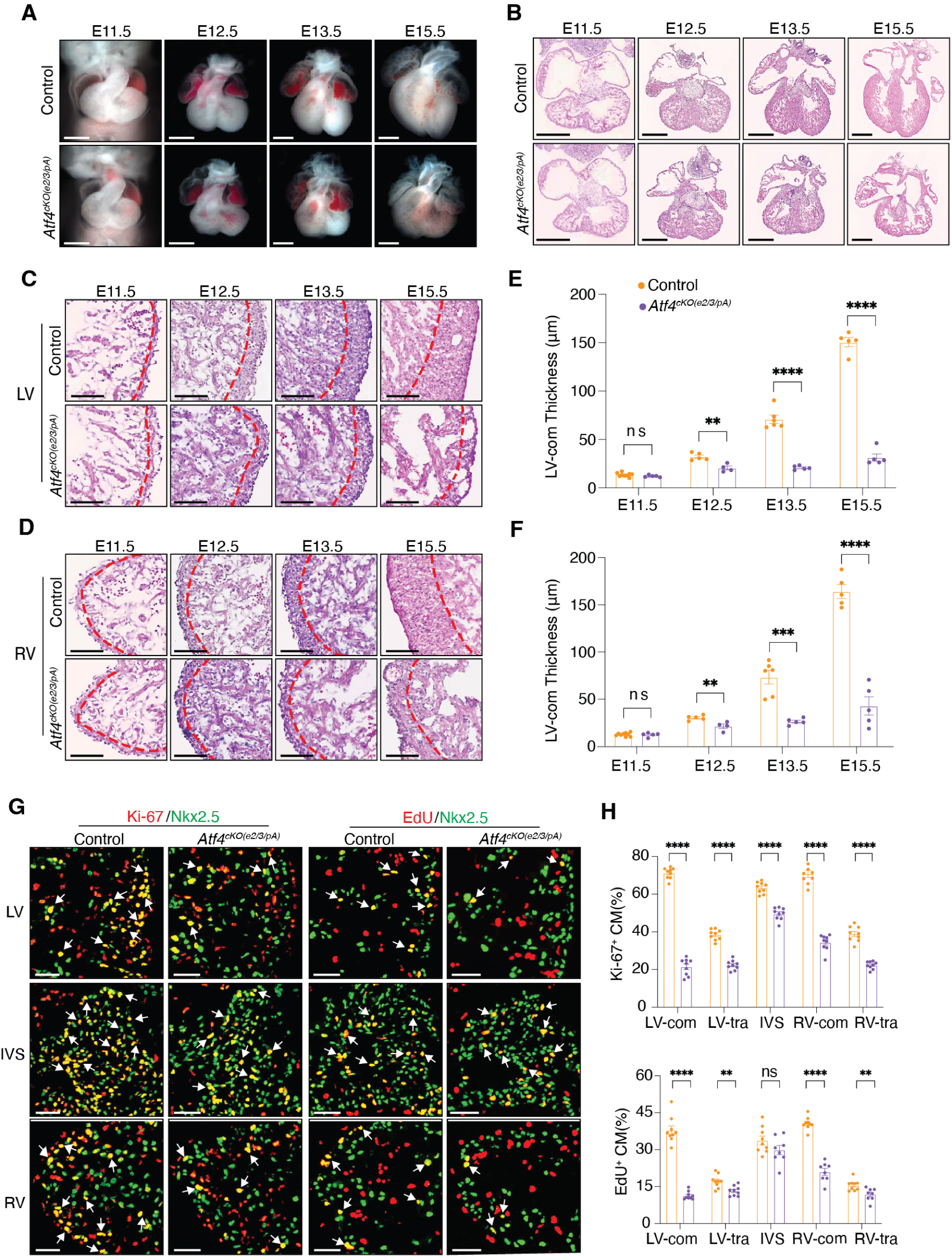
Cardiomyocyte (CM)-specific deletion of *Atf4* results in morphological defects in embryonic hearts. **(A-D)** Representative whole-mount heart **(A)** and H&E stained whole heart **(B)**, left ventricular (LV) **(C)**, and right ventricular (RV) **(D)** images of E11.5 – E15.5 control and *Atf4^cKO(e2/3/pA)^*mouse hearts. The boundary between compact myocardium and trabecular myocardium is depicted by a red dashed line (C-D). Scale bar, 0.5 mm (A-B); 0.1 mm (C-D). **(E-F)** Measurement of thickness of left ventricular compact myocardium (LV-Com) **(E)** and right ventricular compact myocardium (RV-Com) **(F)** thicknesses on control and *Atf4^cKO(e2/3/pA)^*mice heart sections ((n = 4 – 9 hearts per group, n = 4 sections per heart) from E11.5 to E15.5. **(G-H)** Representative immunofluorescent images **(G)** and quantification results **(H)** of Ki-67 positive (Ki-67^+^) and EdU positive CMs on E11.5 control and *Atf4^cKO(e2/3/pA)^* hearts (n = 9 hearts per group, n = 4 - 8 sections per heart). Arrows indicate proliferating CMs (Ki-67^+^/EdU^+^; Nkx2.5^+^). Scale bars, 10 μm. Data are represented as mean±SEM. Statistical significance was determined with 2-tailed Student *t* test (ns, not significant; **P<0.01, ****P<0.0001).

These finding were independently confirmed using a second CM-specific Cre driver, *cTnT*-Cre line^40^, which produced identical cardiac defects (Figure S2D-F). In summary, these results suggested that CM-specific deletion of *Atf4* exons 2-3 leads to defective myocardial growth and embryonic lethality, suggesting a critical role for ATF4 in the developing heart.

### Abnormal activation of p53 pathway in *Atf4^cKO(e2/3/pA)^* mice

To investigate underlying molecular mechanisms by which ATF4 is required for heart development, we performed RNA sequencing (RNA-seq) on ventricular tissues isolated from *Atf4^cKO(e2/3/pA)^* and littermate controls at E11.5. Using false discovery rate (FDR) < 0.05 and fold change ≥ 1.5, we identified 328 differentially expressed genes (DEGs) in *Atf4^cKO(e2/3/pA)^*hearts, with 227 genes significantly upregulated and 101 genes significantly downregulated (Figure 2A). Gene ontology (GO) analysis revealed that p53 signaling was the most enriched pathway in upregulated DEGs (Figure 2B). Gene set enrichment analysis (GSEA) further demonstrated upregulation of p53-related genes in *Atf4^cKO(e2/3/pA)^*hearts (Figure 2C). We confirmed the upregulation of selected p53 downstream key target genes in *Atf4^cKO(e2/3/pA)^* by quantitative real time PCR (qRT-PCR) (Figure S3).

**Figure 2.**
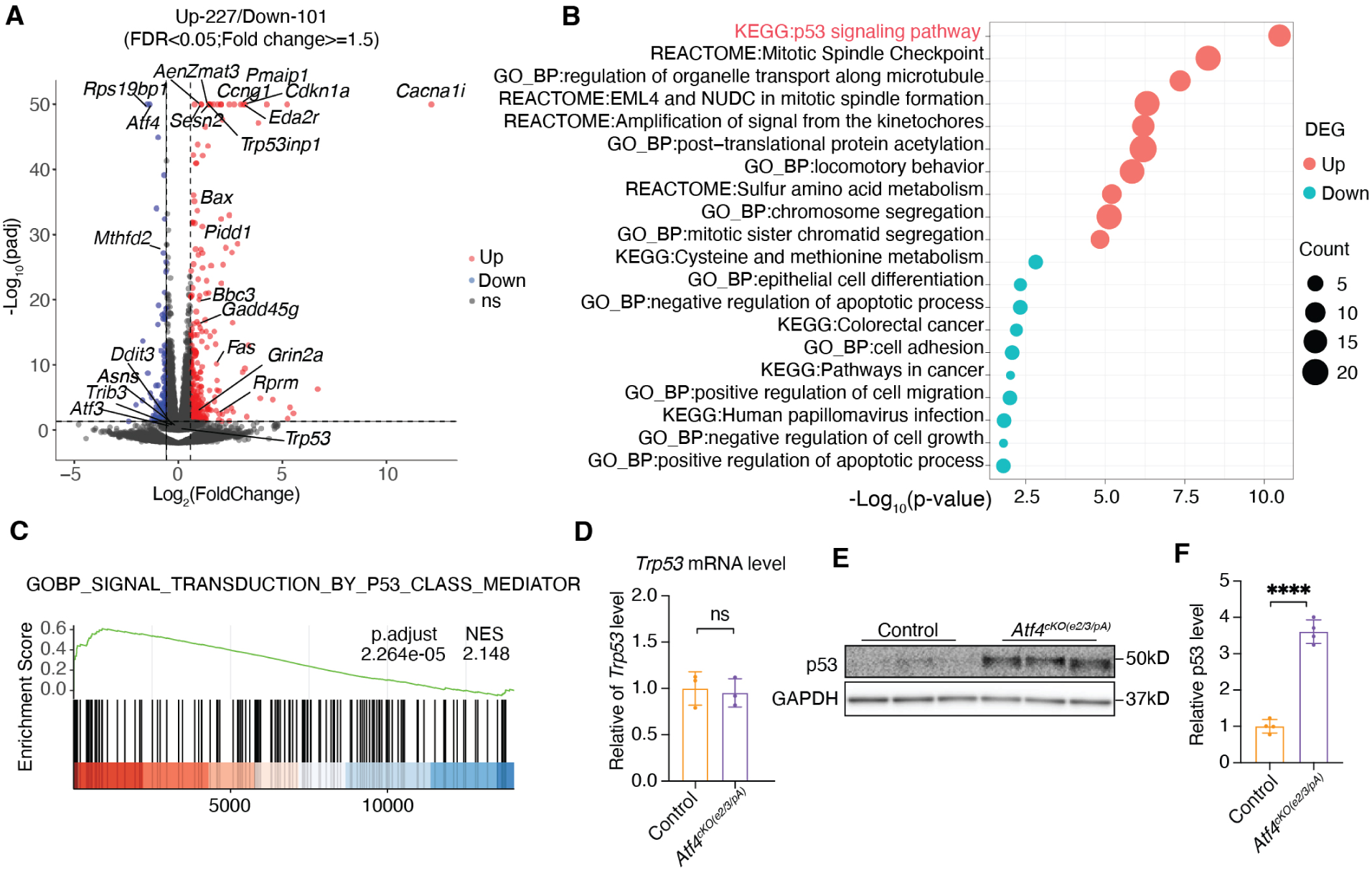
p53 signaling pathway is activated in *Atf4^cKO(e2/3/pA))^* mice. **(A)** Volcano plot of differential expressed genes (DEGs) between control and *Atf4^cKO(e2/3/pA)^* ventricles at E11.5 (n = 4 hearts per group). DEGs with adjusted P<0.05 and fold change ≥ 1.5 are considered significantly upregulated (red dots) or downregulated (blue dots). DEGs with –log_10_(adjusted P values) >50 were limited to 50 in the plot. **(B)** Gene ontology (GO) analysis of upregulated DEGs of *Atf4^cKO(e2/3/pA)^* at E11.5. **(C)** Gene set enrichment analysis (GSEA) of genes in the p53 signaling pathway between control and E11.5 *Atf4^cKO(e2/3/pA)^* heart. **(D)** qRT-qPCR analysis of *Trp53* in control and *Atf4^cKO(e2/3/pA)^* hearts (n = 3 hearts per group) at E11.5. **(E-F)** Western blot **(E)** and accompanying quantitative analysis **(F)** of p53 protein in E11.5 *Atf4^cKO(e2/3/pA)^*(n = 4 hearts per group) compared with littermate control. GAPDH was used as a loading control. Data are represented as mean±SEM. Statistical significance was determined with 2-tailed Student *t* test (ns, not significant; ***P<0.001).

To determine whether upregulation of p53 is responsible for abnormal activation of the p53 pathway in *Atf4^cKO(e2/3/pA)^* hearts, we tested *Trp53* expression levels and p53 protein levels and found that the protein levels of p53 were significantly increased in *Atf4^cKO(e2/3/pA)^*hearts compared to controls (Figure 2E-F), while the mRNA levels of *Trp53,* encoding p53, remained unchanged (Figure 2D). These suggest that *p53* was upregulated in *Atf4^cKO(e2/3/pA)^*hearts through post-transcriptional mechanisms. Given the widely recognized role of the p53 signaling pathway in inhibiting cell proliferation across various tissues^41^, activation of this pathway is likely a significant factor in the reduced CM proliferation seen in *Atf4^cKO(e2/3/pA)^*hearts.

### Transcriptional readthrough of Atf4 leads to the formation of an *Atf4-Cacna1i* fusion transcript and *Rps19bp1* downregulation

Our data indicated that ablating ATF4 in developing CMs led to elevated *p53* protein levels, potentially through post-transcriptional mechanisms. *Rps19bp1*, encoding Active Regulator of SIRT1 (AROS), was among the most downregulated genes in *Atf4^cko(e2/3/pA)^* hearts (Figure 2A and Figure 3A). Previous studies have found that *Rps19bp1* promotes SIRT1-mediated deacetylation of the K382 residue of human p53 (equivalent to K379 in mouse), reducing stability of p53 protein and p53-mediated transcriptional activity^29, 42^. Strikingly, *Rps19bp1* and *Cacna1i*, another gene that was among the most upregulated in the Atf4*^cKO(e2/3/pA)^*heart, are located downstream of *Atf4* transcript in the genome (Figure 3A-B). It raises the possibility that dysregulation of *Rps19bp1* and *Cacna1i* might not be a direct consequence of ATF4 protein loss. Instead, these alterations might stem from disruption of the local transcriptional environment surrounding the *Atf4* locus. Consistent with this, only one canonical ATF4 target gene, *Mthfd2*, showed a mild change in expression, while others, including *Ddit3, Asns, Trib3* and *Atf3*^17, 43–46^, remained unchanged in *Atf4^cKO(e2/3/pA)^* hearts (Figure 2A).

**Figure 3.**
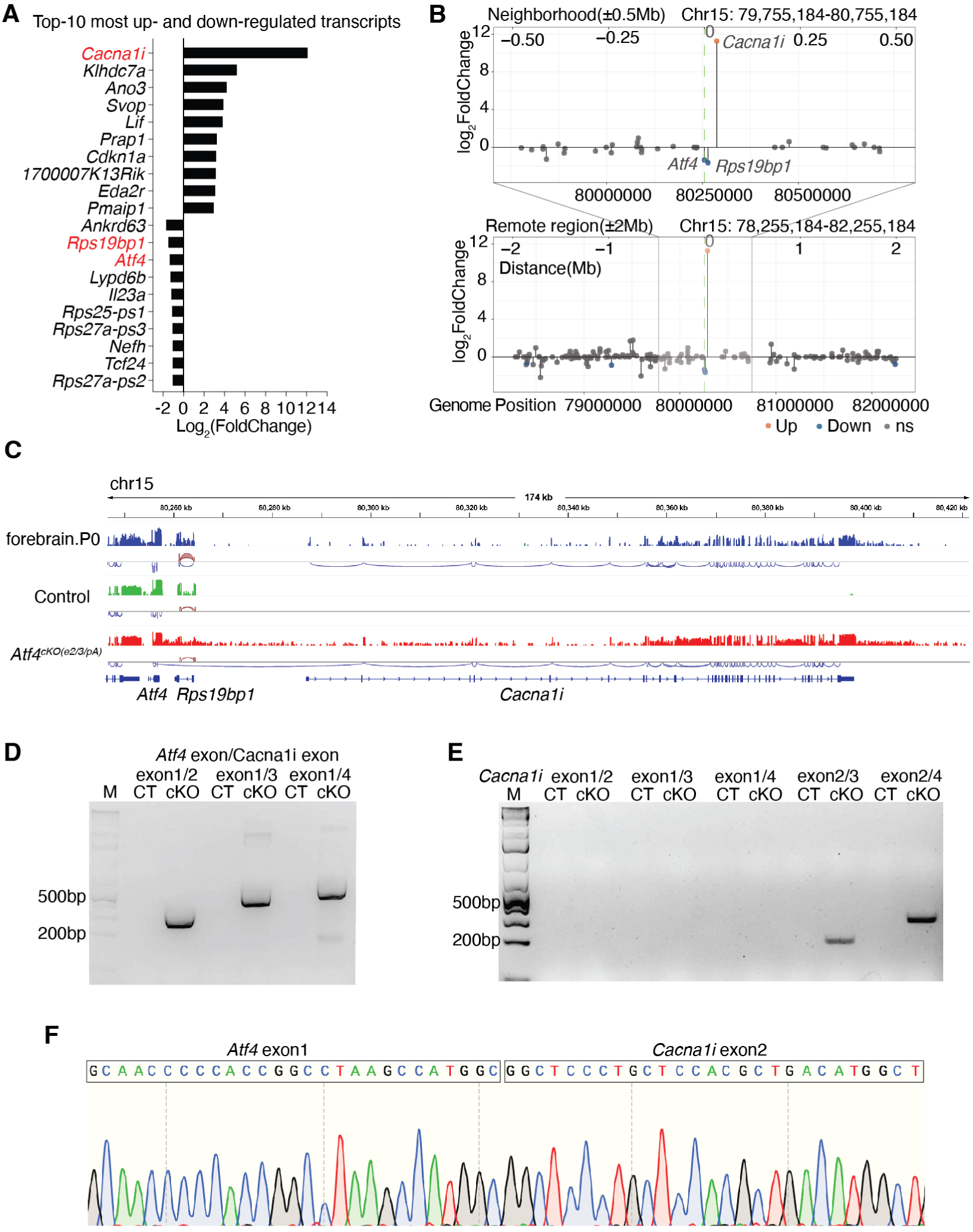
Transcriptional readthrough of *Atf4* leads to the formation of *Atf4*-*Cacna1i* fusion transcript and local transcriptional dysregulation. **(A)** Top-10 most upregulated or downregulated DEGs in *Atf4^cKO(e2/3/pA)^* heart. **(B)** Lollipop plots showing expression changes of genes proximal (±0.5 Mb, top) or distal (±2 Mb, bottom) to *Atf4* in *Atf4^cKO(e2/3/pA)^* hearts. Locations of *Cacna1i*, *Rps19bp1*, and *Atf4* are indicated. **(C)** Normalized RNA-seq tracks and read count for genes adjacent to *Atf4* in E11.5 heart tissues from *Atf4^cKO(e2/3/pA)^* and control. RNA-sequencing data from wildtype P0 mouse hindbrain (SRA: ERS2492429, BioProject: PRJEB26869) show *Cacna1i* expression and splicing under normal conditions. Arcs represent splicing junctions, with red for positive strand junctions and blue for negative strand junctions. **(D-E)** qRT-PCR validation of *Atf4*-*Cacna1i* fusion transcript in from E11.5 control and *Atf4^cKO(e2/3/pA)^* hearts using primers spanning junctions between *Atf4* exon 1 and *Cacna1i* exons 2–4 **(D)**, and within *Cacna1i* (exons 1–4) **(E)**. M: DNA ladder, CT: control, cKO: *Atf4^cKO^*. **(F)** The presence of *Atf4*-*Cacna1i* fusion transcript in *Atf4^cKO^* heart was validated by Sanger sequencing.

We next re-examined our RNA-seq data to identify transcriptomic irregularities within a 1 Mb region centered on *Atf4*. Notably, only *Rps19bp1* and *Cacna1i* were significantly dysregulated within this region, both of which are positioned immediately downstream of *Atf4* (Figure 3B). Interestingly, RNA-seq reads revealed broadly extended RNA coverage spanning the downstream region of *Atf4*, extending as far as 30 kb beyond its canonical transcription termination site (Figure 3C). Furthermore, we found that exon 1 of *Atf4* was aberrantly spliced to exon 2 of *Cacna1i*, resulting in the formation of an *Atf4*-*Cacna1i* fusion transcript in *Atf4^cKO(e2/3/pA)^*samples (Figure 3C), which was further confirmed by qRT-PCR and Sanger sequencing, using primers designed to amplify various exon junctions (Figure 3D-F). Thus, apparent upregulation of *Cacna1i* was owing to misassignment of *Atf4*-*Cacna1i* fusion transcripts as *Cacna1i* transcripts in our initial analyses. The formation of *Atf4*-*Cacna1i* fusion transcripts provides evidence for continued transcription of *Atf4* into the *Cacna1i* locus, a phenomenon known as transcriptional readthrough ^30–32^, likely caused by the deletion of the *Atf4* transcription termination signal in exon 3 of *Atf4*. Notably, *Rps19bp1* is also located downstream of the Atf4 locus, positioned between *Atf4* and *Cacna1i*, but it is transcribed from the opposite strand (Figure 3C). As a result, the transcription complex transcribing the *Atf4*-*Cacna1i* fusion transcript, which is driven by the *Atf4* promoter, may physically collide with the transcription complex transcribing *Rps19bp1*, leading to *Rps19bp1* transcriptional pausing or termination and ultimately *Rps19bp1* downregulation, an example of transcriptional collision^47, 48^ ^30–34, 49^.

To test the hypothesis that transcriptional readthrough caused by deletion of the transcription termination signal in exon 3 underlies the observed phenotypes, we generated a new floxed allele of *Atf4* (*Atf4^fl(e2)/fl(e2)^*) targeting only exon 2 (Figure S5A-B). This design aimed to eliminate ATF4 protein while retaining the endogenous polyA signal located in exon 3 of *Atf4*. We crossed *Atf4^fl(e2)/fl(e2)^* mice with global deleter *Sox2*-cre mice to produce global *Atf4* KO mice (*Atf4^gKO(e2)^*), and with *Xmlc2-Cre* mice to generate CM-specific *Atf4* KO mice (*Atf4^fl(e2)/fl(e2)^*; *Xmlc2-Cre^+/-^*, hereafter *Atf4^cKO(e2)^*). Unexpectedly, both *Atf4^gKO(e2)^* and *Atf4^cKO(e2)^* survived to adulthood without overt abnormalities (Figure S5C-D). In *Atf4^cKO(e2)^* mice, expression of *Rps19bp1* and *Cacna1i* remained unchanged, and normal cardiac function were preserved (Figure S5E-J, Table S2). Transcriptomic profiling by RNA-seq revealed an upregulation of several canonical ATF4 target genes (*Asns*, *Mthfd2*, *Trib3*, etc) in *Atf4^cKO(e2)^* hearts, with GO term analysis highlighting the enrichment of pathways related to amino acid biosynthesis (Figure S5K-L). These results suggested an unexpected increase in ATF4 transcriptional activity. Western blot analysis revealed the presence of a truncated ATF4 protein lacking approximately the first 80 amino acids of its N-terminus (Figure S5M), indicating that *Atf4^cKO(e2)^* mice retain a partial ATF4 product that may retain its transcription factor function. Functional domains located within the N-terminus have been shown to be critical for the transcriptional regulatory function of ATF4 ^50–52^. However, our findings indicated that deletion of the N-terminal region did not abolish ATF4 function *in vivo*. Taken together, these results suggest that exon 2 deletion is insufficient to fully disrupt ATF4 function and thus does not constitute an appropriate model for comprehensive functional analysis of ATF4.

### Loss of ATF4 protein does not impair heart development or function

To further determine whether cardiac defects observed in *Atf4^cKO(e2/3/pA)^* were indeed independent of ATF4 protein function, we generated and characterized additional models ablating ATF4 with minimal genomic perturbation—particularly without disrupting the polyA site—to exclude transcriptional readthrough and its cis-regulatory impact on gene expression. We generated *Atf4^7del/7del^* and *Atf4^1ins/1ins^*mouse models by introducing a small 7bp deletion or a 1bp insertion, respectively, into the coding region of exon 3 (Figure 4A). The indels were introduced into the 5’ end of exon 3 to disrupt the majority of ATF4 coding sequence (Figure 4A). Western blot analyses confirmed a complete loss of ATF4 protein in both models (Figure 4B-C). Remarkably, both mutant lines exhibited normal cardiac morphology and ventricular wall thickness during embryonic development (Figure 4D-G). Although most mutants died perinatally, echocardiography analysis in surviving *Atf4^7del/7del^* and *Atf4^1ins/1ins^*mice showed normal cardiac function (Figure 4H-I, Figure S6A-D, Table S2). RNA-seq analysis of E11.5 *Atf4^7de/7del^* hearts revealed minimal transcriptional changes compared to control hearts, with only 9 DEGs identified—substantially fewer than the 328 DEGs observed in *Atf4^cKO(e2/3/pA)^*hearts, and there was no overlap in the DEG sets between the two mutants (Figure 4J-M). Notably, expression of *Rps19bp1* and *Cacna1i*, as well as p53 target genes remained unaltered (Figure 4G) and only one canonical ATF4 target gene (*Cars*) was mildly downregulated in *Atf4^7de/7del^*mice. Taken together, these findings indicated that ATF4 is dispensable for embryonic cardiac development under physiological conditions.

**Figure 4.**
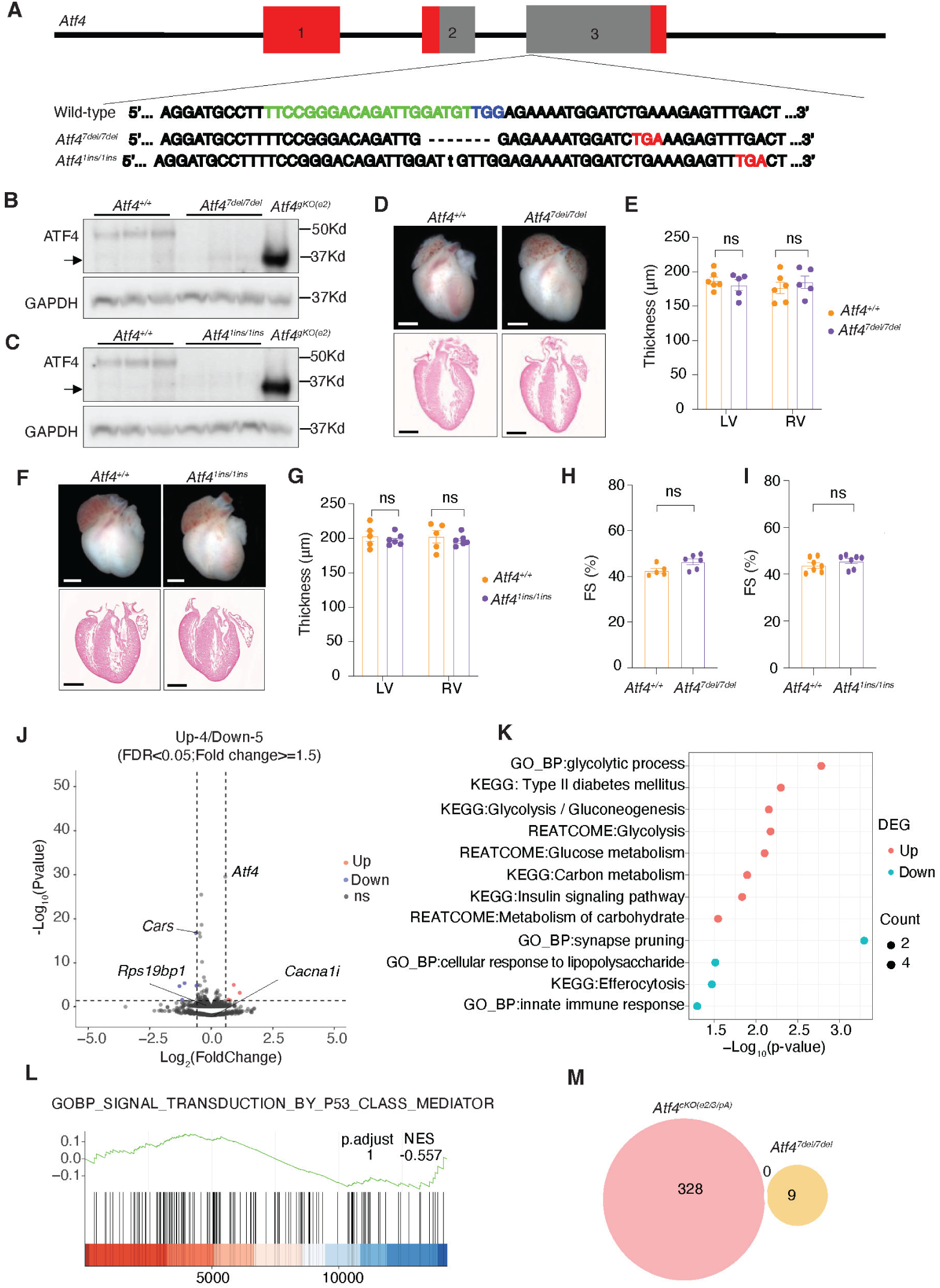
Loss of ATF4 protein does not impair heart development or function. **(A)** Schematic illustrating the generation of the *Atf4* mutant alleles by targeting exon 3 with non-homologous end joining (NHEJ) of CRISPR/Cas9. crRNA sequences are shown in green; predicated premature stop codons are indicated in red. Coding sequences of ATF4 in exon 2 and exon 3 are indicated by grey boxes. **(B-C)** Western blot analysis of ATF4 protein in the large intestine tissue of *Atf4^7del/7del^*and *Atf4^1ins/1ins^* adult mice (n = 3 hearts per group) compared to littermate controls. Arrow indicates truncated ATF4 protein observed in the *Atf4^gKO(e2)^* sample. GAPDH was used as a loading control. **(D-G)** Representative images of whole-mount heart and H&E stained whole heart **(D, F)** and quantification of LV and RV thicknesses **(E, G)** from E17.5 control and *Atf4^7del/7del^* or *Atf4^1ins/1ins^*mouse hearts. Scale bar, 0.5 mm. n = 5 - 6 hearts per group. n = 5 - 6 sections per heart. **(H-I)** Echocardiographic measurements of fractional shortening (FS) in 2-months-old control and *Atf4^7del/7del^*or *Atf4^1ins/1ins^* (n=4-7 mice per group) mice. **(J)** Volcano plot showing differential expressed genes (DEGs) between control and *Atf4^7del/7del^*ventricles at E11.5 (n=4 hearts per group). Significantly upregulated and downregulated genes (adjusted P<0.05 and fold change ≥ 1.5) are marked in red and blue, respectively. **(K)** Gene ontology (GO) analysis of upregulated and downregulated DEGs in *Atf4^7del/7del^* hearts. **(L)** Gene set enrichment analysis (GSEA) of the p53 signaling pathway gene set in E11.5 *Atf4^7del/7del^* heart. **(M)** Venn diagram showing the 0 overlap of DEGs between *Atf4^cKO(e2/3/pA)^* and *Atf4^7del/7del^*hearts. Data are represented as mean±SEM. Statistical significance was determined with 2-tailed Student *t* test (ns, not significant; ***P<0.001).

### Cardiomyocyte-specific deletion of *Rps19bp1* recapitulates the phenotypic and molecular defects of *Atf4^cKO(e2/3/pA)^* mice hearts

The cardiac abnormalities observed in *Atf4^cKO(e2/3/pA)^* mice may result from ectopic p53 pathway activation, which coincided with a marked reduction in *Rps19bp1* expression. We hypothesized that the downregulation of *Rps19bp1* might have contributed to the observed cardiac phenotypes in *Atf4^cKO(e2/3/pA)^*mice. To test this, we crossed *Rps19bp1^fl/fl^* mice with global deleter *Sox2*-cre mice to generate global *Rps19bp1* KO mice (*Rps19bp1^gKO^*), which exhibited embryonic lethality prior to E9.5 (Figure S7A). We then generated CM-specific *Rps19bp1* knockout (KO) mice (*Rps19bp1^fl/fl^*; *Xmlc2-Cre^+/-^*, hereafter *Rps19bp1^cKO^*) using *Xmlc2-Cre*. Consistent with *Atf4^cKO(e2/3/pA)^* mice, *Rps19bp1^cKO^*embryos exhibited abnormal heart morphology at E15.5 and died perinatally (Figure 5A, Figure S7B). Histological analysis revealed a significantly thinner compact myocardium in both ventricles at E15.5 (Figure 5B). To investigate molecular mechanisms underlying these phenotypes, RNA-seq analysis was performed on E11.5 *Rps19bp1^cKO^* hearts. We identified 229 differentially expressed genes (DEGs) (FDR < 0.05, fold change ≥ 1.5) in *Rps19bp1^cKO^* hearts, with 172 genes significantly upregulated and 57 genes significantly downregulated (Figure 5C). Gene ontology (GO) analysis revealed significant enrichment of the p53 signaling pathway in upregulated DEGs (Figure 5D), and Gene set enrichment analysis (GSEA) further demonstrated upregulation of p53-related gene signatures (Figure 5E). Western blotting analysis also revealed significantly increased p53 protein levels in *Rps19bp1^cKO^* hearts compared to controls (Figure 5F). We next compared transcriptional profiles between *Atf4^cKO(e2/3/pA)^* and *Rps19bp1^cKO^* hearts. Remarkably, 108 DEGs were shared between the two models, with a concordance rate of 100% in the direction of expression changes (Figure 5G). Notably, the shared DEGs were again enriched for p53 signaling, supporting a common regulatory mechanism (Figure 5H).

**Figure 5.**
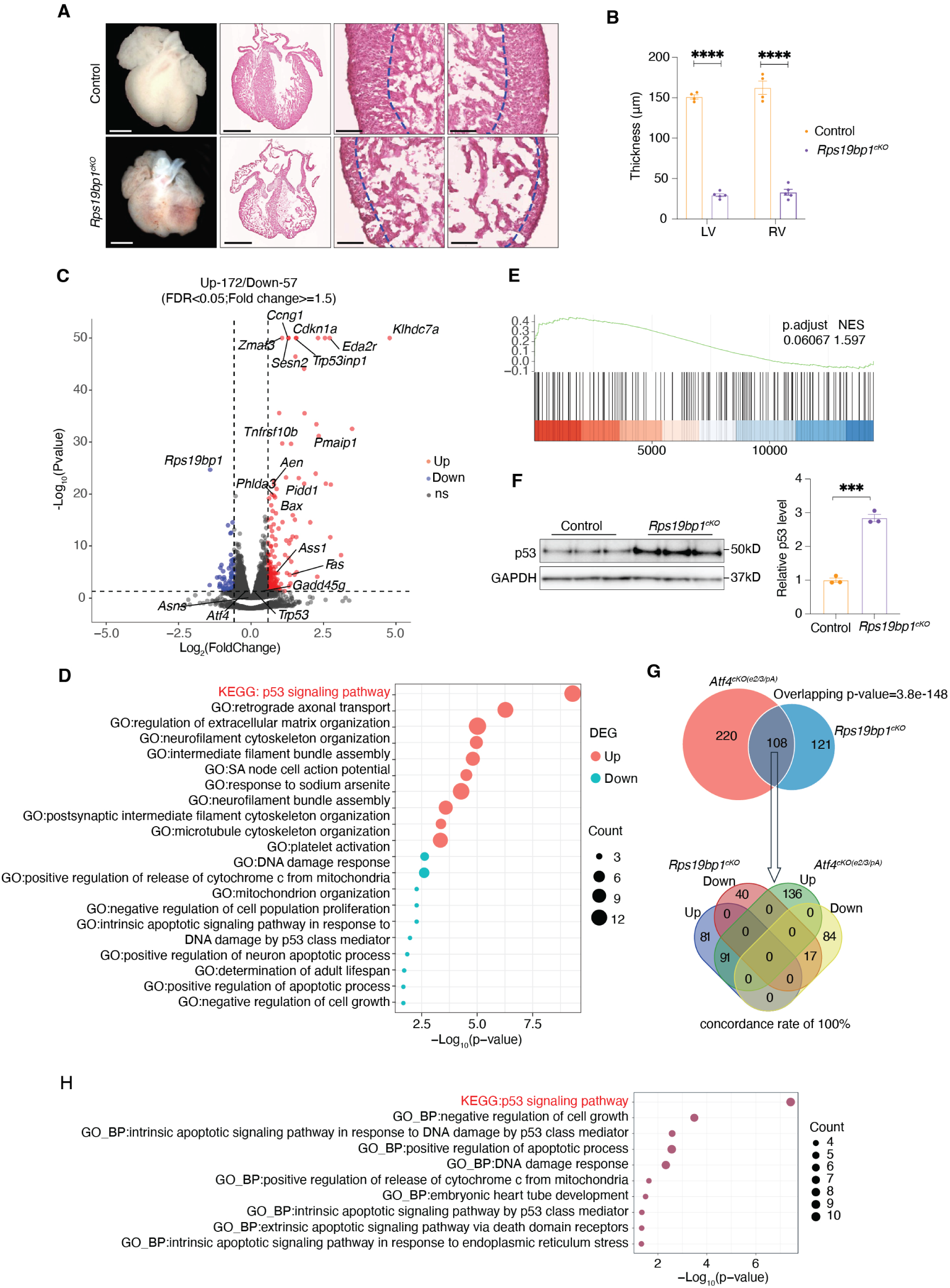
*Rps19bp1^cKO^* mice recapitulate cardiac defects observed in *Atf4^cKO(e2/3/pA)^* mice. **(A)** Representative whole-mount and H&E stained sections of E15.5 hearts from control and *Rps19bp1^cKO^*embryos, including whole heart, LV, and RV. The boundary between compact myocardium and trabecular myocardium is depicted by a blue dashed line. Scale bar, 0.5 mm (whole heart) and 0.1 mm (LV/RV). **(B)** Quantification of LV and RV compact myocardium thicknesses in E15.5 control and *Rps19bp1^cKO^*mice heart (n = 4 - 5 hearts per group; n = 3 sections per heart). **(C)** Volcano plot showing differential expressed genes (DEGs) between control and *Rps19bp1^cKO^*ventricles at E11.5 (n = 4 hearts per group). Significantly upregulated (red) and downregulated (blue) genes were defined as adjusted P<0.05 and fold change ≥ 1.5. DEGs with –log_10_(adjusted P values) >50 were limited to 50 in the plot. **(D)** Gene ontology (GO) analysis of upregulated and downregulated DEGs in *Rps19bp1^cKO^*hearts. **(E)** Gene set enrichment analysis (GSEA) of the p53 signaling pathway in E11.5 *Rps19bp1^cKO^* heart. **(F)** Western blot and accompanying quantitative analysis of p53 protein in E11.5 *Rps19bp1^cKO^* hearts compared with littermate control (n = 3 hearts per group). GAPDH serves as a loading control. **(G)** Venn diagram showing the overlap of 180 DEGs between *Atf4^cKO(e2/3/pA)^* and *Rps19bp1^cKO^* hearts, with a concordance rate of 100%. **(H)** GO analysis of the overlapping DEGs between *Atf4^cKO(e2/3/pA)^*and *Rps19bp1^cKO^* hearts. Data are represented as mean±SEM. Statistical significance was determined with 2-tailed Student *t* test (ns, not significant; ***P<0.001).

These findings demonstrated that *Rps19bp1^cKO^* recapitulated cardiac morphological defects, embryonic lethality, and p53-associated transcriptional changes observed in *Atf4^cKO(e2/3/pA)^* mice (Figure 6). Together, our results establish that downregulation of *Rps19bp1*, rather than the loss of ATF4 function, is the primary driver of abnormal cardiac development in *Atf4^cKO(e2/3/pA)^* hearts.

**Figure 6.**
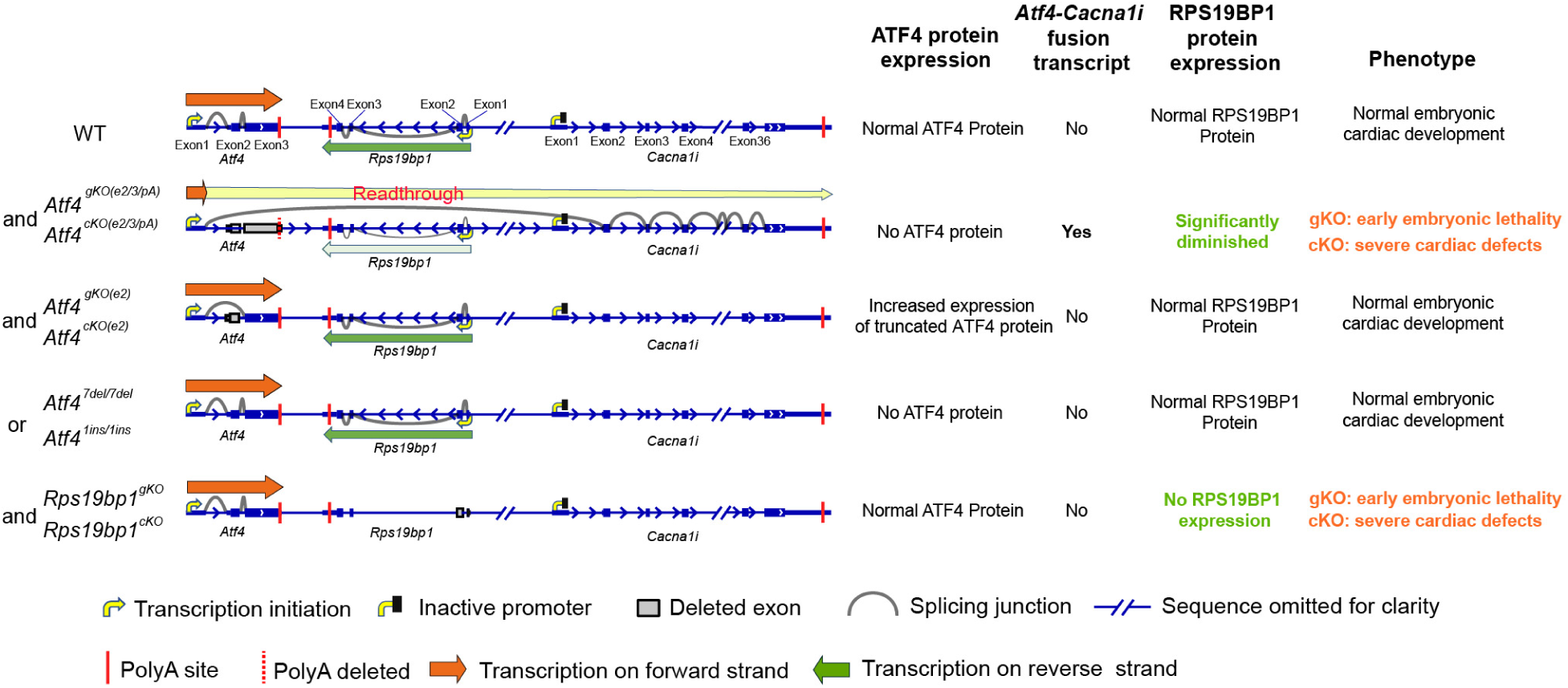
Schematic model illustrating the impact of different *Atf4* and *Rps19bp1* alleles on transcription, protein expression, and cardiac phenotypes. In wild-type (WT) hearts, *Atf4* and its downstream neighbor *Rps19bp1* are independently transcribed, resulting in normal ATF4 and RPS19BP1 protein levels and normal cardiac development. In the widely used *Atf4* model, deletion of exons 2–3 along with the polyA abolishes ATF4 protein production but induces transcriptional readthrough from *Atf4* into *Rps19bp1* and *Cacna1i*. This causes significant repression of *Rps19bp1* and fusion transcript formation (*Atf4*-*Cacna1i*), leading to p53 activation and embryonic lethality in gKO mutants, and severe cardiac defects in cKO mutants. In *Atf4^cKO^ ^(e2)^* allele, deletion of exon 2 leads to significantly increased expression of a truncated ATF4 protein while preserving *Rps19bp1* expression and normal cardiac development. In contrast, frameshift *Atf4* alleles, either *Atf4^7del/7del^* or *Atf4^1ins/1ins^*, abolish ATF4 protein without affecting *Rps19bp1* expression or cardiac development. Finally, CM-specific deletion of *Rps19bp1* alone phenocopies the cardiac defects of *Atf4*, confirming that these phenotypes are driven by transcriptional readthrough-mediated repression rather than ATF4 loss.

## Discussion

In this study, we generated *Atf4* CM-specific KO mice (*Atf4^cKO(e2/3/pA)^*) to study the function of ATF4 in CMs by crossing a widely used *Atf4* floxed mouse line with two CM specific Cre lines. *Atf4^cKO(e2/3/pA)^* mice exhibited abnormal heart morphology, reduced CM proliferation, and embryonic lethality, suggesting that ATF4 plays an important role in developing CMs. However, recently reported CM-specific *Atf4* KO mice generated by *Myh6-Cre* and the same *Atf4* floxed mouse line did not show any basal phenotypes ^15^. This marked phenotypic discrepancy could be due to the relatively low efficiency of *Myh6-Cre* in early embryonic CMs ^37, 53^ compared with the *Xmlc2-Cre* used in our study. At the molecular level, we observed abnormal activation of p53 signaling pathways in *Atf4^cKO(e2/3/pA)^* mice, which might explain the observed reduced CM proliferation and impaired heart development. To investigate the cause of p53 activation and molecular mechanisms underlying cardiac phenotypes in *Atf4^cKO(e2/3/pA)^* mice, we found that *Rps19bp1*, a known p53 suppressor, was drastically downregulated in *Atf4^cKO(e2/3/pA)^*mice. We noticed that *Atf4* and *Rps19bp1* are located within a very short genomic distance. Furthermore, *Cacna1i*, the most upregulated gene in *Atf4^cKO(e2/3/pA)^* mice, is also located proximally to *Atf4*. These observations implied that the dysregulated expression of *Rps19bp1* and *Cacna1i* could be due to genetic disruption at the *Atf4* locus rather than the loss of ATF4 function (Figure 6). This notion is corroborated by findings that *Rps19bp1* and *Cacna1i* were changed similarly in *Atf4*-KO hepatocytes ^54^, hematopoietic stem cell ^28^, skeletal muscle myocytes ^9^ and lens ^55^, all of which utilized the same exons 2–3 floxed *Atf4* allele. Notably, knocking-down or overexpressing ATF4 in hepatocytes had no impact on expression of these two genes ^54^, further suggesting that their dysregulation is independent of ATF4 function.

Our RNA-seq data revealed that upregulation of *Cacna1i* was due to transcriptional readthrough of *Atf4*, forming an *Atf4*-*Cacna1i* fusion transcript, which was improperly interpreted as a marked increase in *Cacna1i* expression. In fact, transcriptional readthrough leading to gene fusion and de novo intergenic splicing has been well documented ^30–32^. The endogenous expression of CACNA1I is not detectable in control hearts (Figure S4). Although the *Atf4*-*Cacna1i* fusion transcript may encode a functional CACNA1I protein—since exon 1 of *Atf4* is non-coding and exon 2 of *Cacna1i* corresponds to its first coding exon—even with the observed increase in transcript levels in the *Atf4^cKO(e2/3/pA)^* heart samples, we were unable to detect any CACNA1I protein (Figure S4). These findings suggest the observed *Cacna1i* upregulation is a consequence of *Atf4* transcriptional readthrough and is unlikely to produce functional proteins to have biological significance in CMs. The potential impact of ectopic expression of neural-specific *Cacna1i* in CMs on cardiac function remain to be determined.

We constructed a new *Atf4* floxed mouse line (*Atf4^fl/fl^*) targeting the first coding exon of *Atf4* (exon 2) to avoid deleting its polyA signal. However, *Atf4^cKO(e2)^* mice produced a truncated ATF4 protein that may partially retain its function. To address this limitation, we generated an *Atf4^7del7/del^* mouse model and an *Atf4^1ins/1ins^* mice model by introducing a small 7bp deletion and a 1bp insertion, respectively, in exon3 of the *Atf4* gene. Neither *Atf4^7del7/del^*nor *Atf4^1ins/1ins^* mice produced truncated ATF4 protein, and both mutants exhibited normal cardiac morphology, with surviving mutants showing normal cardiac function in adult. Furthermore, mRNA expression of *Rps19bp1* and *Cacna1i*, and mRNA and protein levels of p53 remain unchanged in *Atf4^7del/7del^* mice hearts.

*Rps19bp1* is located between *Atf4* and *Cacna1i* and is transcribed from the opposite strand, implying that its downregulation may result from *Atf4* promoter-driven readthrough, as it is well-established that convergent transcription of two bi-directional genes can interfere with the transcription of each other ^30–34^. As a result, the transcription complex transcribing the *Atf4*-*Cacna1i* fusion transcript, which is driven by the strong *Atf4* promoter^56, 57^, may physically collide with the transcription complex transcribing *Rps19bp1*, leading to *Rps19bp1* transcriptional pausing or termination and ultimately *Rps19bp1* downregulation^33, 34^. The significant downregulation of *Rps19bp1* is likely a major contributor to the observed cardiac phenotypes in *Atf4^cKO(e2/3/pA)^*mice. Indeed, *Rps19bp1^cKO^* mice displayed identical cardiac morphological defects, embryonic lethality, and molecular defects to those observed in *Atf4^cKO(e2/3/pA)^*mice (Figure 6). This further supports the conclusion that the downregulation of *Rps19bp1*, rather than loss of ATF4 function, contributes to the abnormal cardiac development.

As a ubiquitously-expressed protein, RPS19BP1 may also play important roles in other tissues. Significant fold-change reductions in *Rps19bp1* expression have also been noted in *Atf4* KOs generated using the widely used floxed allele of *Atf4* exon 2 and 3 plus polyA signal in hepatocytes ^54^, hematopoietic stem cells ^28^, and skeletal muscle myocytes ^9^. Additionally, other *Atf4* null allele mice, generated by replacing exon 2 and exon 3 plus polyA signal with a neomycin phosphotransferase cassette, albeit exhibiting distinct phenotypes including defects in lens development and severe fetal anemia ^22, 23^, also showed *Rps19bp1* downregulation ^55^, and p53 ablation rescued lens defects in one of the lines of *Atf4* null mice ^22^. Importantly, however, adult skeletal muscle fibers do not exhibit a basal phenotype or increased expression of p53 when *Atf4* is deleted using *MCK-Cre* and the same floxed *Atf4* mice line, although there is a reduction in *Atf4* and *Rps19bp1* levels ^9^. This may be because the absolute change in *Rps19bp1* expression is minimal, given that its normal expression in adult skeletal muscle fibers is already low ^9^. Furthermore, when *Atf4* KO skeletal muscle fibers are exposed to atrophy-inducing stress conditions, they manifest a phenotype (decreased muscle atrophy) that is recapitulated by RNAi targeting *Atf4* and strongly accentuated by concomitant p53 ablation ^9–12^. Conversely, forced expression of ATF4 in adult skeletal muscle fibers induces the opposite phenotype (increased muscle atrophy), which is enhanced by co-expression of p53 ^10, 11^.

Thus, in some contexts, changes in *Rps19bp1* expression due to excision of *Atf4* exons 2 and 3 and the *Atf4* polyA sequence appear to have little if any phenotypic effects, which may be related to context-specific differences in basal *Rps19bp1* expression. However, in other contexts, including but perhaps not limited to the developing heart, the only biologically important consequence of deleting *Atf4* exons 2 and 3 and polyA sequences is likely due to the reduction in *Rps19bp1* expression. Based on this new finding (Figure 6), it now seems critically important to recognize that any phenotype observed following excision of *Atf4* exons 2 and 3 and polyA sequence could be potentially caused by loss of ATF4 and/or loss of p53 inhibition by RPS19BP1, and therefore, both possibilities should be rigorously evaluated by multiple complementary approaches.

In summary, we provided unequivocal evidence that ATF4 itself is not essential for heart development and adult function and that loss of *Rps19bp1*, not *Atf4*, in cardiomyocytes lead to abnormal cardiac development (Figure 6). Our findings also revealed that *Atf4* transcriptional readthrough, rather than loss of ATF4 function, results in the dysregulation of *Rps19bp1*, underlying the phenotypic and molecular changes in *Atf4^cKO(e2/3/pA)^*mice. All previous *Atf4* knockout studies employed similar strategies of deleting exons 2/3 or exon 3 plus polyA of the *Atf4* gene ^19, 22, 23, 28, 58–60^, which can lead to *Atf4* transcriptional readthrough. Consistently, *Rps19bp1* downregulation has been observed across various tissues in multiple *Atf4* global or tissue-specific knockout models, suggesting that the phenotypic and molecular changes in previous studies may likely result from *Atf4* readthrough-induced *Rps19bp1* downregulation. Our study underscores the importance of re-evaluating previous *Atf4* findings and highlights the need for caution in future research to avoid potential misinterpretations of results. Furthermore, it emphasizes the necessity of considering potential “neighborhood effects” on gene expression, which can significantly impact the interpretation of the data obtained in gene knockout experiments.

## Supporting information

Figure S1-7; Table S1-2; Full unedited gels

## Acknowledgments

1. Z. Zhang, T. Wu, Z. Chen, D. Chen, S. Evans, X. Zhou, and J. Chen designed the research. Z. Zhang, T. Wu, Z. Chen, Z. Liang, Y. Gu, M. Ye, and F. Barroga performed the research. Z. Zhang, T. Wu, Z. Chen, D. Chen, Z. Liang, S. Evans, X. Zhou, and J. Chen analyzed the data. Z. Zhang, T. Wu, Z. Chen, D. Chen, S. Evans, X. Zhou, and J. Chen wrote the article. C. Adams provided the *Atf4^fl(e2/3/pA)/fl(e2/3/pA)^* mouse line and edited the manuscript.

## Supplementary Materials

Figures S1 to S7

Table S1 to S2 Full unedited gels

## References

1. Adams CM, Ebert SM, Dyle MC. Role of ATF4 in skeletal muscle atrophy. Curr Opin Clin Nutr Metab Care 2017;20:164–168.

2. Ameri K, Harris AL. Activating transcription factor 4. Int J Biochem Cell Biol 2008;40:14–21.

3. Kasai S, Yamazaki H, Tanji K, Engler MJ, Matsumiya T, Itoh K. Role of the ISR-ATF4 pathway and its cross talk with Nrf2 in mitochondrial quality control. J Clin Biochem Nutr 2019;64:1–12.

4. Wortel IMN, van der Meer LT, Kilberg MS, van Leeuwen FN. Surviving Stress: Modulation of ATF4-Mediated Stress Responses in Normal and Malignant Cells. Trends Endocrinol Metab 2017;28:794–806.

5. Li K, Xiao Y, Yu J, Xia T, Liu B, Guo Y, Deng J, Chen S, Wang C, Guo F. Liver-specific Gene Inactivation of the Transcription Factor ATF4 Alleviates Alcoholic Liver Steatosis in Mice. J Biol Chem 2016;291:18536–18546.

6. Zhang Q, Yu J, Liu B, Lv Z, Xia T, Xiao F, Chen S, Guo F. Central activating transcription factor 4 (ATF4) regulates hepatic insulin resistance in mice via S6K1 signaling and the vagus nerve. Diabetes 2013;62:2230–2239.

7. Baleriola J, Walker CA, Jean YY, Crary JF, Troy CM, Nagy PL, Hengst U. Axonally synthesized ATF4 transmits a neurodegenerative signal across brain regions. Cell 2014;158:1159–1172.

8. Rozpedek W, Pytel D, Mucha B, Leszczynska H, Diehl JA, Majsterek I. The Role of the PERK/eIF2α/ATF4/CHOP Signaling Pathway in Tumor Progression During Endoplasmic Reticulum Stress. Curr Mol Med 2016;16:533–544.

9. Miller MJ, Marcotte GR, Basisty N, Wehrfritz C, Ryan ZC, Strub MD, McKeen AT, Stern JI, Nath KA, Rasmussen BB, Judge AR, Schilling B, Ebert SM, Adams CM. The transcription regulator ATF4 is a mediator of skeletal muscle aging. Geroscience 2023;45:2525–2543.

10. Fox DK, Ebert SM, Bongers KS, Dyle MC, Bullard SA, Dierdorff JM, Kunkel SD, Adams CM. p53 and ATF4 mediate distinct and additive pathways to skeletal muscle atrophy during limb immobilization. Am J Physiol Endocrinol Metab 2014;307:E245–261.

11. Ebert SM, Monteys AM, Fox DK, Bongers KS, Shields BE, Malmberg SE, Davidson BL, Suneja M, Adams CM. The transcription factor ATF4 promotes skeletal myofiber atrophy during fasting. Mol Endocrinol 2010;24:790–799.

12. Ebert SM, Dyle MC, Kunkel SD, Bullard SA, Bongers KS, Fox DK, Dierdorff JM, Foster ED, Adams CM. Stress-induced skeletal muscle Gadd45a expression reprograms myonuclei and causes muscle atrophy. J Biol Chem 2012;287:27290–27301.

13. Jeong MH, Jeong HJ, Ahn BY, Pyun JH, Kwon I, Cho H, Kang JS. Correction: PRMT1 suppresses ATF4-mediated endoplasmic reticulum response in cardiomyocytes. Cell Death Dis 2020;11:203.

14. Masuda M, Miyazaki-Anzai S, Keenan AL, Shiozaki Y, Okamura K, Chick WS, Williams K, Zhao X, Rahman SM, Tintut Y, Adams CM, Miyazaki M. Activating transcription factor-4 promotes mineralization in vascular smooth muscle cells. JCI Insight 2016;1:e88646.

15. Wang X, Zhang G, Dasgupta S, Niewold EL, Li C, Li Q, Luo X, Tan L, Ferdous A, Lorenzi PL, Rothermel BA, Gillette TG, Adams CM, Scherer PE, Hill JA, Wang ZV. ATF4 Protects the Heart From Failure by Antagonizing Oxidative Stress. Circ Res 2022;131:91–105.

16. Fu HY, Okada K, Liao Y, Tsukamoto O, Isomura T, Asai M, Sawada T, Okuda K, Asano Y, Sanada S, Asanuma H, Asakura M, Takashima S, Komuro I, Kitakaze M, Minamino T. Ablation of C/EBP homologous protein attenuates endoplasmic reticulum-mediated apoptosis and cardiac dysfunction induced by pressure overload. Circulation 2010;122:361–369.

17. Freundt JK, Frommeyer G, Wötzel F, Huge A, Hoffmeier A, Martens S, Eckardt L, Lange PS. The Transcription Factor ATF4 Promotes Expression of Cell Stress Genes and Cardiomyocyte Death in a Cellular Model of Atrial Fibrillation. Biomed Res Int 2018;2018:3694362.

18. Yao Y, Lu Q, Hu Z, Yu Y, Chen Q, Wang QK. A non-canonical pathway regulates ER stress signaling and blocks ER stress-induced apoptosis and heart failure. Nat Commun 2017;8:133.

19. Masuoka HC, Townes TM. Targeted disruption of the activating transcription factor 4 gene results in severe fetal anemia in mice. Blood 2002;99:736–745.

20. Zhao Y, Zhou J, Liu D, Dong F, Cheng H, Wang W, Pang Y, Wang Y, Mu X, Ni Y, Li Z, Xu H, Hao S, Wang X, Ma S, Wang QF, Xiao G, Yuan W, Liu B, Cheng T. ATF4 plays a pivotal role in the development of functional hematopoietic stem cells in mouse fetal liver. Blood 2015;126:2383–2391.

21. Fischer C, Johnson J, Stillwell B, Conner J, Cerovac Z, Wilson-Rawls J, Rawls A. Activating transcription factor 4 is required for the differentiation of the lamina propria layer of the vas deferens. Biol Reprod 2004;70:371–378.

22. Hettmann T, Barton K, Leiden JM. Microphthalmia due to p53-mediated apoptosis of anterior lens epithelial cells in mice lacking the CREB-2 transcription factor. Dev Biol 2000;222:110–123.

23. Tanaka T, Tsujimura T, Takeda K, Sugihara A, Maekawa A, Terada N, Yoshida N, Akira S. Targeted disruption of ATF4 discloses its essential role in the formation of eye lens fibres. Genes Cells 1998;3:801–810.

24. Yang X, Matsuda K, Bialek P, Jacquot S, Masuoka HC, Schinke T, Li L, Brancorsini S, Sassone-Corsi P, Townes TM, Hanauer A, Karsenty G. ATF4 is a substrate of RSK2 and an essential regulator of osteoblast biology; implication for Coffin-Lowry Syndrome. Cell 2004;117:387–398.

25. Xiao G, Jiang D, Ge C, Zhao Z, Lai Y, Boules H, Phimphilai M, Yang X, Karsenty G, Franceschi RT. Cooperative interactions between activating transcription factor 4 and Runx2/Cbfa1 stimulate osteoblast-specific osteocalcin gene expression. J Biol Chem 2005;280:30689–30696.

26. Zhang Y, Lin T, Lian N, Tao H, Li C, Li L, Yang X. Hop2 Interacts with ATF4 to Promote Osteoblast Differentiation. J Bone Miner Res 2019;34:2287–2300.

27. Dobreva G, Chahrour M, Dautzenberg M, Chirivella L, Kanzler B, Fariñas I, Karsenty G, Grosschedl R. SATB2 is a multifunctional determinant of craniofacial patterning and osteoblast differentiation. Cell 2006;125:971–986.

28. Zheng Z, Yang S, Gou F, Tang C, Zhang Z, Gu Q, Sun G, Jiang P, Wang N, Zhao X, Kang J, Wang Y, He Y, Yang M, Lu T, Lu S, Qian P, Zhu P, Cheng H, Cheng T. The ATF4-RPS19BP1 axis modulates ribosome biogenesis to promote erythropoiesis. Blood 2024;144:742–756.

29. Kim EJ, Kho JH, Kang MR, Um SJ. Active regulator of SIRT1 cooperates with SIRT1 and facilitates suppression of p53 activity. Mol Cell 2007;28:277–290.

30. Osato N, Suzuki Y, Ikeo K, Gojobori T. Transcriptional interferences in cis natural antisense transcripts of humans and mice. Genetics 2007;176:1299–1306.

31. Rutkowski AJ, Erhard F, L’Hernault A, Bonfert T, Schilhabel M, Crump C, Rosenstiel P, Efstathiou S, Zimmer R, Friedel CC, Dölken L. Widespread disruption of host transcription termination in HSV-1 infection. Nat Commun 2015;6:7126.

32. Muniz L, Deb MK, Aguirrebengoa M, Lazorthes S, Trouche D, Nicolas E. Control of Gene Expression in Senescence through Transcriptional Read-Through of Convergent Protein-Coding Genes. Cell Rep 2017;21:2433–2446.

33. Shearwin KE, Callen BP, Egan JB. Transcriptional interference--a crash course. Trends Genet 2005;21:339–345.

34. Wang L, Watters JW, Ju X, Lu G, Liu S. Head-on and co-directional RNA polymerase collisions orchestrate bidirectional transcription termination. Mol Cell 2023;83:1153–1164.e1154.

35. Fang X, Stroud MJ, Ouyang K, Fang L, Zhang J, Dalton ND, Gu Y, Wu T, Peterson KL, Huang HD, Chen J, Wang N. Adipocyte-specific loss of PPAR. JCI Insight 2016;1:e89908.

36. Breckenridge R, Kotecha S, Towers N, Bennett M, Mohun T. Pan-myocardial expression of Cre recombinase throughout mouse development. Genesis 2007;45:135–144.

37. Wu T, Liang Z, Zhang Z, Liu C, Zhang L, Gu Y, Peterson KL, Evans SM, Fu XD, Chen J. PRDM16 Is a Compact Myocardium-Enriched Transcription Factor Required to Maintain Compact Myocardial Cardiomyocyte Identity in Left Ventricle. Circulation 2022;145:586–602.

38. Zhou X, Fang X, Ithychanda SS, Wu T, Gu Y, Chen C, Wang L, Bogomolovas J, Qin J, Chen J. Interaction of Filamin C With Actin Is Essential for Cardiac Development and Function. Circ Res 2023;133:400–411.

39. Liu C, Spinozzi S, Chen JY, Fang X, Feng W, Perkins G, Cattaneo P, Guimarães-Camboa N, Dalton ND, Peterson KL, Wu T, Ouyang K, Fu XD, Evans SM, Chen J. Nexilin Is a New Component of Junctional Membrane Complexes Required for Cardiac T-Tubule Formation. Circulation 2019;140:55–66.

40. Jiao K, Kulessa H, Tompkins K, Zhou Y, Batts L, Baldwin HS, Hogan BL. An essential role of Bmp4 in the atrioventricular septation of the mouse heart. Genes Dev 2003;17:2362–2367.

41. Engeland K. Cell cycle regulation: p53-p21-RB signaling. Cell Death Differ 2022;29:946–960.

42. Knight JR, Allison SJ, Milner J. Active regulator of SIRT1 is required for cancer cell survival but not for SIRT1 activity. Open Biol 2013;3:130130.

43. Fawcett TW, Martindale JL, Guyton KZ, Hai T, Holbrook NJ. Complexes containing activating transcription factor (ATF)/cAMP-responsive-element-binding protein (CREB) interact with the CCAAT/enhancer-binding protein (C/EBP)-ATF composite site to regulate Gadd153 expression during the stress response. Biochem J 1999;339 ( **Pt 1**):135–141.

44. Fusakio ME, Willy JA, Wang Y, Mirek ET, Al Baghdadi RJ, Adams CM, Anthony TG, Wek RC. Transcription factor ATF4 directs basal and stress-induced gene expression in the unfolded protein response and cholesterol metabolism in the liver. Mol Biol Cell 2016;27:1536–1551.

45. Balasubramanian MN, Butterworth EA, Kilberg MS. Asparagine synthetase: regulation by cell stress and involvement in tumor biology. Am J Physiol Endocrinol Metab 2013;304:E789–799.

46. Neill G, Masson GR. A stay of execution: ATF4 regulation and potential outcomes for the integrated stress response. Front Mol Neurosci 2023;16:1112253.

47. Webster MW, Takacs M, Zhu C, Vidmar V, Eduljee A, Abdelkareem M, Weixlbaumer A. Structural basis of transcription-translation coupling and collision in bacteria. Science 2020;369:1355–1359.

48. Prescott EM, Proudfoot NJ. Transcriptional collision between convergent genes in budding yeast. Proc Natl Acad Sci U S A 2002;99:8796–8801.

49. Hobson DJ, Wei W, Steinmetz LM, Svejstrup JQ. RNA polymerase II collision interrupts convergent transcription. Mol Cell 2012;48:365–374.

50. Liang G, Hai T. Characterization of human activating transcription factor 4, a transcriptional activator that interacts with multiple domains of cAMP-responsive element-binding protein (CREB)-binding protein. J Biol Chem 1997;272:24088–24095.

51. Lassot I, Estrabaud E, Emiliani S, Benkirane M, Benarous R, Margottin-Goguet F. p300 modulates ATF4 stability and transcriptional activity independently of its acetyltransferase domain. J Biol Chem 2005;280:41537–41545.

52. Schoch S, Cibelli G, Magin A, Steinmüller L, Thiel G. Modular structure of cAMP response element binding protein 2 (CREB2). Neurochem Int 2001;38:601–608.

53. Chen JW, Zhou B, Yu QC, Shin SJ, Jiao K, Schneider MD, Baldwin HS, Bergelson JM. Cardiomyocyte-specific deletion of the coxsackievirus and adenovirus receptor results in hyperplasia of the embryonic left ventricle and abnormalities of sinuatrial valves. Circ Res 2006;98:923–930.

54. Byles V, Cormerais Y, Kalafut K, Barrera V, Hughes Hallett JE, Sui SH, Asara JM, Adams CM, Hoxhaj G, Ben-Sahra I, Manning BD. Hepatic mTORC1 signaling activates ATF4 as part of its metabolic response to feeding and insulin. Mol Metab 2021;53:101309.

55. Xiang J, Pompetti AJ, Faranda AP, Wang Y, Novo SG, Li DW, Duncan MK. ATF4 May Be Essential for Adaption of the Ocular Lens to Its Avascular Environment. Cells 2023;12.

56. Dey S, Savant S, Teske BF, Hatzoglou M, Calkhoven CF, Wek RC. Transcriptional repression of ATF4 gene by CCAAT/enhancer-binding protein β (C/EBPβ) differentially regulates integrated stress response. J Biol Chem 2012;287:21936–21949.

57. Vattem KM, Wek RC. Reinitiation involving upstream ORFs regulates ATF4 mRNA translation in mammalian cells. Proc Natl Acad Sci U S A 2004;101:11269–11274.

58. Kitakaze K, Oyadomari M, Zhang J, Hamada Y, Takenouchi Y, Tsuboi K, Inagaki M, Tachikawa M, Fujitani Y, Okamoto Y, Oyadomari S. ATF4-mediated transcriptional regulation protects against β-cell loss during endoplasmic reticulum stress in a mouse model. Mol Metab 2021;54:101338.

59. Guo Q, Xu Z, Zhou D, Fu T, Wang W, Sun W, Xiao L, Liu L, Ding C, Yin Y, Zhou Z, Sun Z, Zhu Y, Zhou W, Jia Y, Xue J, Chen Y, Chen XW, Piao HL, Lu B, Gan Z. Mitochondrial proteostasis stress in muscle drives a long-range protective response to alleviate dietary obesity independently of ATF4. Sci Adv 2022;8:eabo0340.

60. Li K, Zhang J, Yu J, Liu B, Guo Y, Deng J, Chen S, Wang C, Guo F. MicroRNA-214 suppresses gluconeogenesis by targeting activating transcriptional factor 4. J Biol Chem 2015;290:8185–8195.

