## Supplementary material for "Transcriptional Readthrough at *Atf4* Locus Suppresses *Rps19bp1* and Impairs Heart Development": Figure S1-7; Table S1-2; Full unedited gels

**Contents:**

1) Figures S1-7 and Figure Legends.
2) Tables S1-2.
3) Full unedited gels.

| | $Atf4^{gko(e2/3/pA)/gko(e2/3/pA)}$ | $Atf4^{gko(e2/3/pA)/+}$ | $Atf4^{+/+}$ |
|---|---|---|---|
| E8.5 | 0 | 29 | 11 |
| E9.5 | 0 | 12 | 9 |

**Figure S1.** Numbers of viable embryos from each genotype ($Atf4^{+/+}$, $Atf4^{gKO(e2/3/pA)/+}$ and $Atf4^{gKO(e2/3/pA)/\ gKO(e2/3/pA)}$) from the crossbreeding of $Atf4^{gKO(e2/3/pA)/+}$ mice.

**A**

| | $Atf4^{fl(e2/3/pA)/fl(e2/3/pA); Cre+}$ | $Atf4^{fl(e2/3/pA)/+; Cre+}$ | $Atf4^{fl(e2/3/pA)/fl(e2/3/pA); Cre-}$ | $Atf4^{fl(e2/3/pA)/+; Cre-}$ |
|---|---|---|---|---|
| E11.5 | 37 | 45(1*) | 43 | 44(1*) |
| E15.5 | 11(1*) | 10 | 6 | 9 |
| E17.5 | 21(17*) | 25 | 34 | 21(1*) |
| P1 | 1(1*) | 19 | 10 | 26 |
| P21 | 0 | 29 | 21 | 36 |

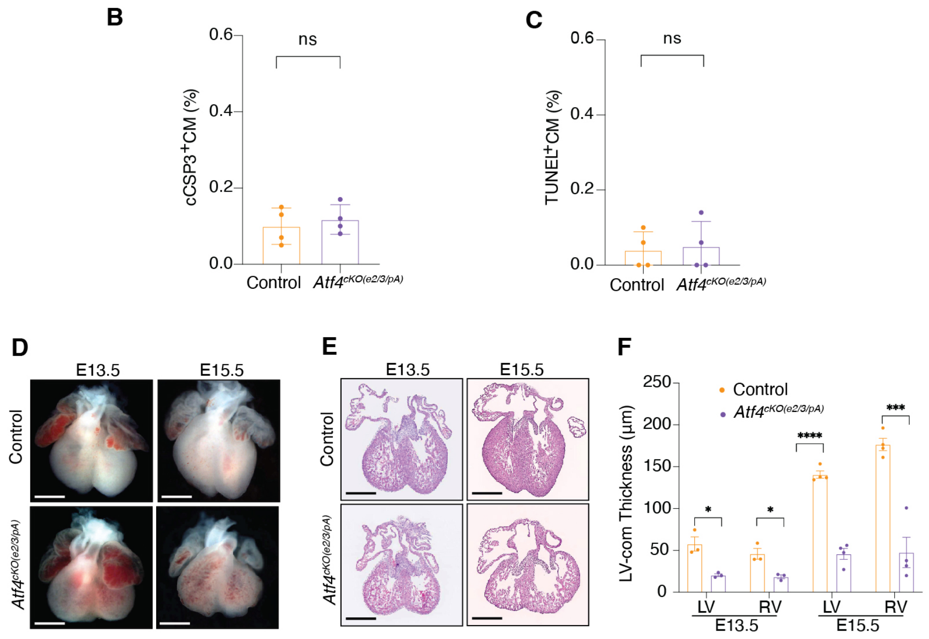


**Figure S2. (A)** Numbers of control ($Atf4^{fl(e2/3/pA)/fl(e2/3/pA)}$or $Atf4^{fl(e2/3/pA)/+}$), $Atf4^{cKO(e2/3/pA)}$ ($Atf4^{fl(e2/3/pA)/fl(e2/3/pA)}$; *Xmlc2-Cre*$^{+/-}$) and $Atf4^{cHet}$ ($Atf4^{fl(e2/3/pA)/+}$; *Xmlc2-Cre*$^{+/-}$) embryos or pups from crossing $Atf4^{fl(e2/3/pA)/+}$); *Xmlc2-Cre*$^{+/-}$ males with $Atf4^{fl(e2/3/pA)/fl(e2/3/pA)}$ females. Data are presented as the total number of embryos or pups, with the number of dead embryos or pups shown in parentheses and indicated by an asterisk (*). **(B-C)** Quantitative analysis of cleaved caspase 3 positive (cCSP3$^{+}$) CMs and TUNEL$^{+}$ CMs from E11.5 control and $Atf4^{cKO(e2/3/pA)}$ hearts. CMs are marked by Nkx2-5. **(D-F)** Representative Whole-mount **(D)** and H&E stained cryosection **(E)** images, and quantitative analysis **(F)** of E13.5 and E15.5 control compared to $Atf4^{cKO(e2/3/pA)}$ ($Atf4^{fl(e2/3/pA)/fl(e2/3/pA)}$; *cTNT-Cre*$^{+/-}$) heart. Scale bar, 0.5mm. Data are represented as mean±SEM. Statistical significance was determined with 2-tailed Student *t* test (ns, not significant; *P<0.05, ***P<0.001, ****P<0.0001).

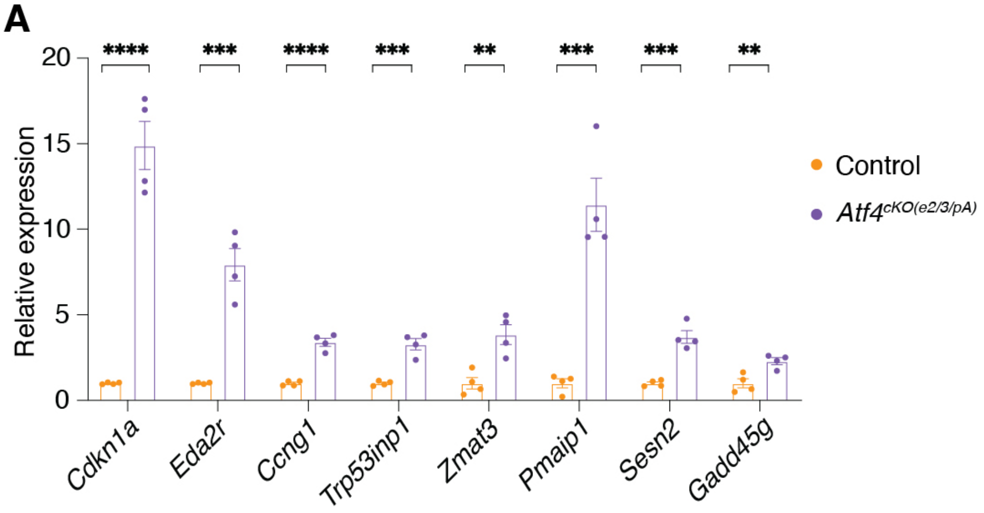


**Figure S3.** qRT-PCR analysis of selected p53 target genes in *Atf4*$^{cKO(e2/3/pA)}$ and control hearts at E11.5. Data are represented as mean±SEM. Statistical significance was determined with 2-tailed Student *t* test (**P<0.01, ***P<0.001, ****P<0.0001).

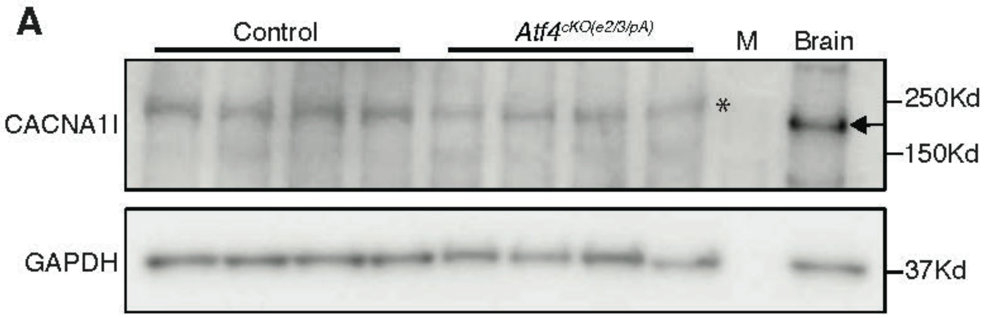


**Figure S4.** Western blot of CACNA1i protein in E11.5 *Atf4*$^{cKO(e2/3/pA)}$ (n = 4 hearts per group) compared with littermate control. M, Protein Marker. Mouse brain sample was used as a positive control. Asterisk (*) denotes a nonspecific band, and the arrow indicates the CACNA1I protein.

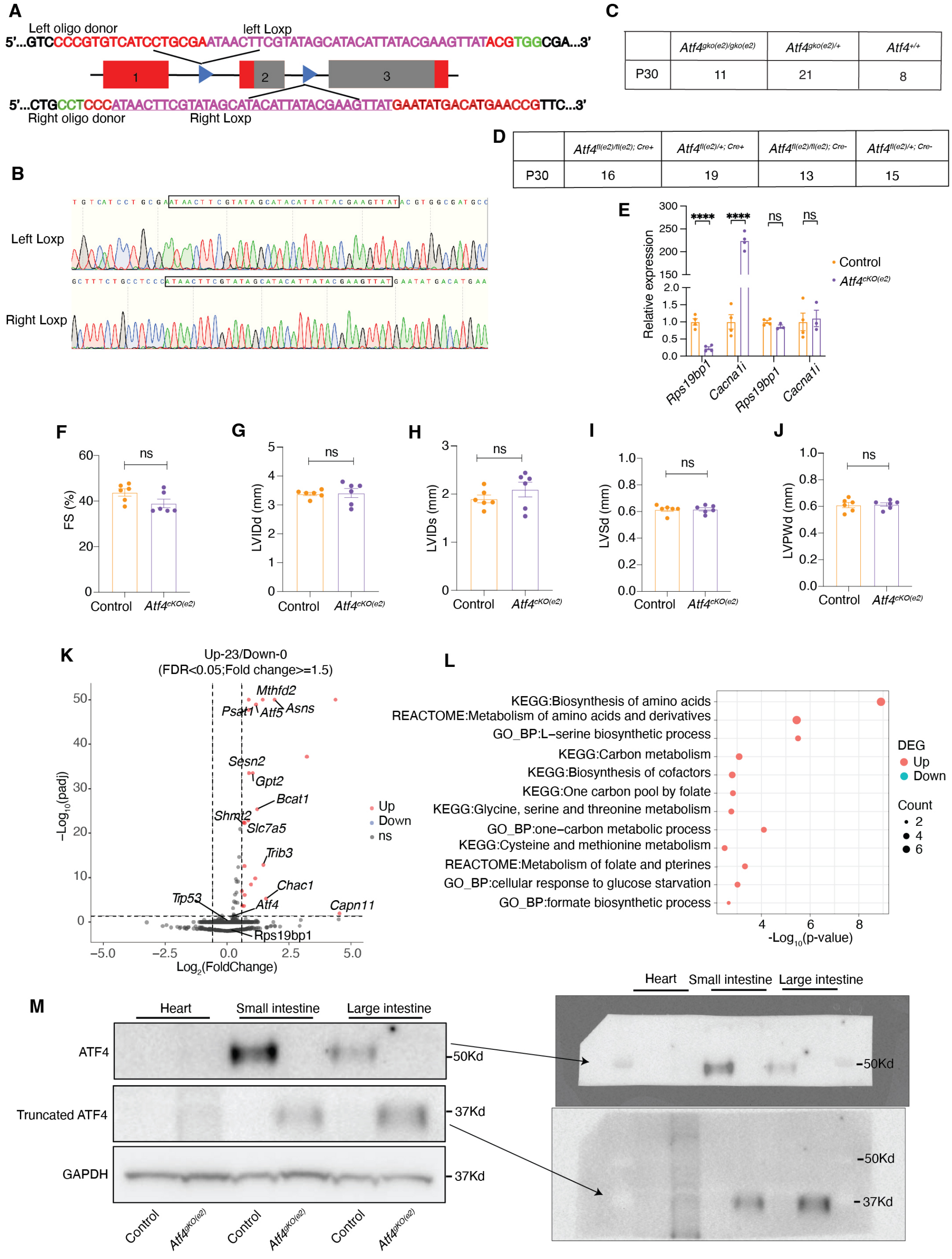


C

| | Atf4gko(e2)/gko(e2) | Atf4gko(e2)/+ | Atf4+/+ |
|---|---|---|---|
| P30 | 11 | 21 | 8 |

D

| | Atf4fl(e2)/fl(e2); Cre+ | Atf4fl(e2)/+; Cre+ | Atf4fl(e2)/fl(e2); Cre- | Atf4fl(e2)/+; Cre- |
|---|---|---|---|---|
| P30 | 16 | 19 | 13 | 15 |

**Figure S5. (A)** Schematic illustration to generate the new *Atf4* floxed allele mice by insertion of two LoxP sites. The sequences of CRISPR RNA (crRNA) to target the Cas9 nuclease to intron 1 (left) and intron 2 (right) are shown in red. The protospacer adjacent motif (PAM) is shown in green. the LoxP site is highlighted in purple in the oligo donor sequence. **(B)** The successful integration of LoxP sites was confirmed by Sanger sequencing. **(C)** Number of viable one-month-old $Atf4^{+/+}$, $Atf4^{(e2)/+}$ and $Atf4^{gKO(e2)/\ gKO(e2)}$ mice. **(D)** Number of viable one-month-old control ($Atf4^{fl(e2)/fl(e2)}$or $Atf4^{fl(e2)/+}$), $Atf4^{cKO(e2)}$ ($Atf4^{fl(e2)/fl(e2)}$; *Xmlc2-Cre*$^{+/-}$) and $Atf4^{cHet}$ ($Atf4^{fl(e2)/+}$); *Xmlc2-Cre*$^{+/-}$) mice from crossing $Atf4^{fl(e2)/+}$); *Xmlc2-Cre*$^{+/-}$ males with $Atf4^{fl(e2)/fl(e2)}$ females. **(E)** Quantitative real-time PCR (qRT-PCR) analysis of mRNA expression level of *Rps19pb1* and *Cacna1i* in control and $Atf4^{cKO(e2)}$ hearts at E11.5. **(F-J)** Echocardiographic measurements of (F) fractional shortening (FS); (G) left ventricle internal diameter, diastole (LVIDd); (H) left ventricle internal diameter, systole (LVIDs); (I) interventricular septum diameter (IVSd); and (J) LV posterior wall, diastole (LVPWd) of 3-month-old control (n=6) and $Atf4^{cKO(e2)}$ (n = 6) mice **(K)** Volcano plot of differential expressed genes (DEGs) between control and $Atf4^{cKO(e2)}$ ventricles at E11.5 (n = 4). DEGs with adjusted $P<0.05$ and fold change $\geq 1.5$ are considered significantly upregulated (red dots) or downregulated genes (blue dots). **(L)** Gene ontology (GO) analysis of upregulated and downregulated DEGs of $Atf4^{cKO(e2)}$ at E11.5. **(M)** Western blot analysis of ATF4 protein in heart, small intestine and large intestine tissues in E11.5 $Atf4^{gKO(e2)}$ mice compared with littermate controls. Notably, the truncated ATF4 protein (~37 kDa) in the mutant tissues is expressed at much higher levels than the full-length wild-type ATF4 protein (~50 kDa) in control tissues. Only the truncated ATF4 protein is detectable with short exposure (bottom right panel), whereas the wild-type ATF4 protein requires prolonged exposure to be visualized (upper right panel). GAPDH was used as a loading control. Data are represented as mean±SEM. Statistical significance was determined with 2-tailed Student t test (ns, not significant; **$P<0.01$, ****$P<0.0001$).

**A**

| Stage | *Atf4*$^{7del/7del}$ | *Atf4*$^{7del/+}$ | *Atf4*$^{+/+}$ |
|---|---|---|---|
| E11.5 | 12 | 30 | 12 |
| E17.5 | 12 | 15 | 15 |
| P0 | 14(11*)(13.6%) | 37(1*) | 23(1*) |
| P7 | 4(13%) | 59 | 32 |
| P21 | 6(11%) | 112 | 54 |

**B**

| Stage | *Atf4*$^{1ins/1ins}$ | *Atf4*$^{1ins/+}$ | *Atf4*$^{+/+}$ |
|---|---|---|---|
| E11.5 | 9 | 39 | 25 |
| E17.5 | 7 | 17 | 5 |
| P0 | 7(3*)(12%) | 42(4*) | 35(2*) |
| P7 | 11(1*)(17%) | 87 | 60 |
| P21 | 14(1*)(18%) | 107 | 72 |

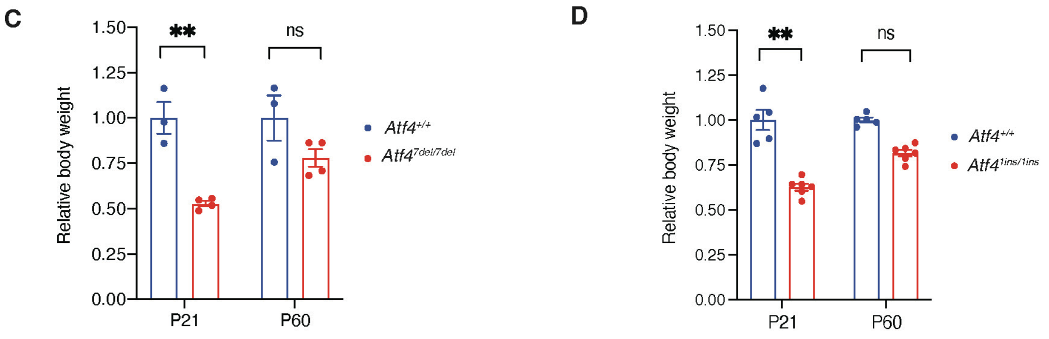


**Figure S6. (A)** Numbers of each genotype (*Atf4*$^{+/+}$, *Atf4*$^{7del/+}$ and *Atf4*$^{7del/7del}$) in survived offsprings from the crossbreeding of *Atf4*$^{7del/+}$ mice. **(B)** Numbers of each genotype (*Atf4*$^{+/+}$, *Atf4*$^{1ins/+}$ and *Atf4*$^{1ins/1ins}$) in survived offsprings from the crossbreeding of *Atf4*$^{1ins/+}$ mice. Data are presented as the total number of pups, with the number of dead pups shown in parentheses and indicated by an asterisk (*), and the proportion of surviving *Atf4*$^{7del/7del}$ or *Atf4*$^{1ins/1ins}$ pups relative to *Atf4*$^{+/+}$ shown in parentheses as a percentage. **(C-D)** Relative body weight of *Atf4*$^{+/+}$, *Atf4*$^{7del/7del}$ or *Atf4*$^{1ins/1ins}$ mice at postnatal day 21 (P21) and day 60 (P60). Data are represented as mean±SEM. Statistical significance was determined with 2-tailed Student t test (ns, not significant; **P<0.01, ****P<0.0001).

**A**

| Stage | *Rps19bp1*$^{-/-}$ | *Rps19bp1*$^{+/-}$ | *Rps19bp1*$^{+/+}$ |
|---|---|---|---|
| E8.5 | 4 | 17 | 11 |
| E9.5 | 2(2*) | 25 | 11 |

**B**

| | *Rps19bp1*$^{fl/fl;\ Cre+}$ | *Rps19bp1*$^{fl/fl;\ Cre-}$ | *Rps19bp1*$^{fl/+;\ Cre+}$ | *Rps19bp1*$^{fl/+;\ Cre-}$ |
|---|---|---|---|---|
| E11.5 | 22 | 22 | 15 | 16 |
| E15.5 | 7 | 11 | 21 | 27 |
| E17.5 | 9(3*) | 13 | 10 | 13 |
| E18.5 | 9(6*) | 13 | 13(2*) | 17 |
| P0 | 2(2*) | 21(2*) | 16(1*) | 14(2*) |
| P1 | 0 | 19 | 15(1*) | 12 |
| P21 | 0 | 18 | 15(1*) | 11 |

**Figure S7. (A)** Numbers of each genotype (*Rps19bp1*$^{+/+}$*, Rps19bp*$^{+/-}$ and *Rps19bp1*$^{-/-}$ ) in survived offsprings from the crossbreeding of *Rps19bp1*$^{+/-}$ mice. **(B)** Numbers of control (*Rps19bp1*$^{fl/fl}$ or *Rps19bp1*$^{fl/+}$), *Rps19bp1*$^{cKO}$ (*Rps19bp1*$^{fl/fl}$; *Xmlc2-Cre*$^{+/-}$) and *Rps19bp1*$^{cHet}$ (*Rps19bp1*$^{fl/+}$; *Xmlc2-Cre*$^{+/-}$) from crossing *Rps19bp1*$^{fl/+}$; *Xmlc2-Cre*$^{+/-}$ males with *Rps19bp1*$^{fl/fl}$ females. Data are presented as the total number of embryos or pups, with the number of dead embryos or pups shown in parentheses and indicated by an asterisk (*).

**Table S1.** Primers information.

| Name | Sequences | |
|---|---|---|
| *Atf4-fl(e2/3)* -P1 | GCGGAGGAGTGTCTAAAGCGCTAC | *Atf4* exon2/3 floxed mice |
| *Atf4-fl(e2/3)* -P2 | GGTTGCACAAGATGGAGGCTTAGC | |
| *Atf4-fl(e2/3)* -P3 | GCAGACGTTCCTGGGTTAGA | |
| *Atf4-fl(e2/3)* -P4 | GCTTCCTGCCTACATTGCTC | |
| *Xml*-Cre-F | AGCCATCTTTGGTTCTCTGC | for *Xmcl2-Cre* |
| *Xml*-Cre-R | TCCCTGAACATGTCCATCAG | |
| Cre-long-P1: | GAGCATACCTGGAAAATGCTTC | for *Sox2-cre* |
| Cre-Long-P2: | CCGGCAAAACAGGTAGTTATTC | |
| *Atf4-fl(e2)* -P1 | GTCTAAAGCGCTACTGCTGC | Atf4 exon2 floxed mice |
| *Atf4-fl(e2)* -P2 | GCTTGTCCTCGCGTCCTAATA | |
| *Atf4-fl(e2)* -P3 | GGAATGGCCGGCTATGGAT | |
| *Atf4-fl(e2)* -P4 | CAACGTGGTCAAGAGCTCATC | |
| mouse sry (sex) F | AACAACTGGGCTTTGCACATTG | for mouse sex |
| mouse sry (sex) R | GTTTATCAGGGTTTCTCTCTAGC | |
| *trp53* F | GAGACCGCCGTACAGAAGAA | for qRT-PCR |
| *trp53* R | ATCTCGAAGCGTTTACGCCC | |
| 18S F | GGAAGGGCACCACCAGGAGT | |
| 18S R | TGCAGCCCCGGACATCTAAG | |
| *Cdkn1a* F | TGTCGCTGTCTTGCACTCTG | |
| *Cdkn1a* R | GTGGGCACTTCAGGGTTTTC | |
| *Eda2r* F | TGTTGCTACACGTGGAAGTGA | |
| *Eda2r* R | GGCCACATCTACTTATCGCTCT | |
| *Ccng1* F | AGACGTGGCTGTCAAGATGAT | |
| *Ccng1* R | AGTTTCAAGCCGCAGACCTT | |
| *Trp53inp1* F | AGACTCACGGGCACAGAAATG | |
| Trp53inp1 R | GGGCGAAAACTCTTGGGTTG | |
| *Zmat3* F | GTCACCTTGAACTCCGCTCA | |
| *Zmat3* R | GGACAGCTGTTAGCTGCGTA | |
| *Sesn2* F | TGCCATTCCGAGATCAAGGG | |
| *Sesn2* R | GGGTGTAGACCCATCACCAC | |
| *Pmaip1* F | CTGAGATGCCCGGGAGAAAG | |
| *Pmaip1* R | TGCGCCAGAACCACAGTTAT | |
| *Gadd45g* F | CGTCTACGAGTCCGCCAAA | |
| *Gadd45g* R | CACGCGCACGATATCAATGT | |
| *Atf4* F | CCTATAAAGGCTTGCGGCCA | |
| *Atf4* R | GATTTCGTGAAGAGCGCCAT | |
| *Rps19bp1* F | GAAGGGAAAGGCGCCTAAGT | |
| *Rps19bp1* R | CTTTTCGGCCTTGGTTCTGC | |
| *Cacna1i* F | GTCTTCACCAAGATGGACGACC | |
| *Cacna1i* R | ACTTCGCACCAGTCAGGCTTGT | |
| *Atf4*-wt-tF1 | CGGCACACGCGGTTTTACAAGC | *Atf4* exon2/3 floxed mice without polyA site |
| *Atf4*-wt-tR1 | AATCCCAGAAGCGTCTACAGGG | |
| *Atf4*-wt-tF2 | TGACCCACCTGGAGTTAGTTTG | |
| *Atf4*-wt-tR2 | TCCCTCATCTTCACCGTCTCA | |
| *Cacna1i* e12 F | TAAGGACGGCTTTTCCTCGC | *Cacna1i* exon1 and exon2 primers for qPCR |
| *Cacna1i* e12 R | GACGTCAGGGTTGGTTCCAT | |
| *Cacna1i* e13 F | TCCCGGAATCTGAGGCGT | *Cacna1i* exon1 and exon3 primers for qPCR |
| *Cacna1i* e13 R | GCATGCTCACACACTCGAAC | |
| *Cacna1i* e14 F | ATAAGGACGGCTTTTCCTCGC | *Cacna1i* exon1 and exon4 primers for qPCR |
| *Cacna1i* e14 R | CCCCGAGGTAGCACTTCTTG | |
| *Atf4* exon1 F | GCTGTCTGCCGGTTTAAGTTG | *Atf4* exon1 and *Cacna1i* junction primer for qPCR |
| *Cacna1i* exon2 F | CTGACGTCCCGCATCCAGAC | *Cacna1i* exon2 or exon3 or exon4 primers for qPCR |
| *Cacna1i* exon3 R | GCAGGATCTTGCAGCGGTCT | |
| *Cacna1i* exon4 R | CGGTTCCATGTGTCCCCGAG | |
| *Atf4e3* wt F12 | TTCCGGGACAGATTGGATG | *Atf4 7del/7del* and *1ins/1ins* PCR dection |
| *Atf4e3* 1ins F14 | TTCCGGGACAGATTGGATTG | |
| *Atf4e3* 7del F14 | TTTCCGGGACAGATTGGAGA | |
| *Atf4e3* del F14 | AAGGGTGAGTAGGACGGCT | *Atf4* exon3 mutations sequencing |
| *Atf4e3* del R14 | TTGGGTTCACTGTCTGAGGG | |
| *Rps19bp1*-tF1 | AGCCTTGAACATGGTTAGGCAAC | *Rps19bp1* exon2/3 floxed mice |
| *Rps19bp1*-tR1 | AAGTACCCTCAGGCACAGAGTAAGC | |
| *Rps19bp1*-tF2 | GCATCGCATTGTCTGAGTAGGTG | |
| *Rps19bp1*-tR2 | CCTGAGCAGAGCACATTAAGAGC | |

**Table S2.** Echocardiographic parameters.

| Genotype | BW(g) | Sex | HR M-mode (bpm) | LVIDd(mm) | LVIDs(mm) | %FS | IVSd(mm) | LVPWd(mm) | LVIDd/LVPWd | LVIDd/BW | LVM(d)(mg) | LVM/BW (mg/g) |
|---|---|---|---|---|---|---|---|---|---|---|---|---|
| | 20.0 | M | 650 | 3.22 | 1.73 | 46.4 | 0.63 | 0.66 | 4.85 | 0.161 | 61.47 | 3.07 |
| | 24.0 | M | 665 | 3.35 | 1.91 | 43.2 | 0.80 | 0.78 | 4.28 | 0.140 | 87.12 | 3.63 |
| *Atf4*$^{+/+}$ | 25.0 | M | 669 | 3.49 | 2.04 | 41.7 | 0.78 | 0.74 | 4.72 | 0.140 | 87.84 | 3.51 |
| | 19.0 | F | 690 | 3.42 | 2.04 | 40.5 | 0.68 | 0.67 | 5.11 | 0.180 | 72.16 | 3.80 |
| | 24.0 | M | 656 | 3.57 | 2.11 | 41.0 | 0.69 | 0.66 | 5.43 | 0.149 | 76.68 | 3.20 |
| | 18.0 | M | 639 | 3.17 | 1.64 | 48.4 | 0.67 | 0.68 | 4.70 | 0.176 | 63.20 | 3.51 |
| | 18.0 | M | 658 | 2.88 | 1.53 | 46.9 | 0.68 | 0.68 | 4.27 | 0.160 | 54.54 | 3.03 |
| *Atf4*$^{7del/7del}$ | 20.0 | M | 659 | 3.16 | 1.80 | 43.1 | 0.72 | 0.72 | 4.37 | 0.158 | 69.56 | 3.48 |
| | 20.0 | M | 677 | 3.17 | 1.66 | 47.5 | 0.82 | 0.77 | 4.13 | 0.159 | 79.29 | 3.96 |
| | 14.0 | F | 691 | 2.94 | 1.48 | 49.8 | 0.65 | 0.68 | 4.31 | 0.210 | 55.21 | 3.94 |
| | 16.0 | M | 676 | 2.71 | 1.57 | 42.0 | 0.67 | 0.68 | 3.99 | 0.169 | 49.08 | 3.07 |

| Genotype | BW(g) | Sex | HR M-mode (bpm) | LVIDd(mm) | LVIDs(mm) | %FS | IVSd(mm) | LVPWd(mm) | LVIDd/LVPWd | LVIDd/BW | LVM(d)(mg) | LVM/BW (mg/g) |
|---|---|---|---|---|---|---|---|---|---|---|---|---|
| | 24.0 | M | 656 | 3.67 | 1.91 | 47.9 | 0.76 | 0.72 | 5.07 | 0.153 | 91.41 | 3.81 |
| | 23.0 | M | 700 | 3.51 | 1.84 | 47.5 | 0.69 | 0.68 | 5.19 | 0.152 | 75.79 | 3.30 |
| | 23.0 | M | 626 | 3.61 | 2.16 | 40.2 | 0.70 | 0.68 | 5.31 | 0.157 | 80.98 | 3.52 |
| *Atf4*$^{+/+}$ | 22.0 | M | 635 | 3.58 | 1.97 | 44.9 | 0.65 | 0.70 | 5.10 | 0.163 | 78.11 | 3.55 |
| | 18.0 | F | 644 | 3.09 | 1.75 | 43.4 | 0.69 | 0.70 | 4.43 | 0.172 | 63.30 | 3.52 |
| | 20.0 | F | 684 | 3.51 | 2.08 | 40.8 | 0.66 | 0.68 | 5.16 | 0.176 | 74.37 | 3.72 |
| | 22.6 | M | 645 | 3.38 | 1.97 | 41.9 | 0.66 | 0.64 | 5.31 | 0.150 | 66.87 | 2.96 |
| | 19.1 | M | 656 | 3.08 | 1.79 | 41.9 | 0.66 | 0.64 | 4.79 | 0.161 | 57.61 | 3.02 |
| | 19.3 | M | 608 | 3.14 | 1.70 | 46.0 | 0.76 | 0.70 | 4.51 | 0.163 | 69.71 | 3.61 |
| | 18.7 | M | 681 | 3.20 | 1.69 | 47.3 | 0.64 | 0.68 | 4.74 | 0.171 | 62.00 | 3.32 |
| *Atf4*$^{1ins/1ins}$ | 18.0 | M | 619 | 3.03 | 1.60 | 47.3 | 0.68 | 0.67 | 4.52 | 0.168 | 58.93 | 3.27 |
| | 17.0 | F | 655 | 2.97 | 1.75 | 41.1 | 0.68 | 0.70 | 4.23 | 0.175 | 59.10 | 3.48 |
| | 20.0 | M | 657 | 2.81 | 1.49 | 47.0 | 0.72 | 0.76 | 3.70 | 0.140 | 59.54 | 2.98 |
| | 17.0 | F | 674 | 2.90 | 1.50 | 48.2 | 0.68 | 0.66 | 4.38 | 0.170 | 54.26 | 3.19 |

| Genotype | BW(g) | Sex | HR M-mode (bpm) | LVIDd(mm) | LVIDs(mm) | %FS | IVSd(mm) | LVPWd(mm) | LVIDd/LVPWd | LVIDd/BW | LVM(d)(mg) | LVM/BW (mg/g) |
|---|---|---|---|---|---|---|---|---|---|---|---|---|
| | 21.0 | F | 694 | 3.33 | 1.82 | 45.4 | 0.62 | 0.61 | 5.45 | 0.159 | 60.99 | 2.90 |
| | 21.0 | F | 639 | 3.41 | 1.88 | 44.7 | 0.63 | 0.54 | 6.29 | 0.162 | 58.94 | 2.81 |
| Control | 20.0 | F | 705 | 3.14 | 1.64 | 47.7 | 0.55 | 0.57 | 5.52 | 0.157 | 48.52 | 2.43 |
| | 23.0 | M | 688 | 3.54 | 2.21 | 37.6 | 0.62 | 0.63 | 5.65 | 0.154 | 68.19 | 2.96 |
| | 29.0 | M | 671 | 3.41 | 2.03 | 40.5 | 0.62 | 0.67 | 5.10 | 0.118 | 67.56 | 2.33 |
| | 22.0 | F | 699 | 3.45 | 1.83 | 46.8 | 0.65 | 0.64 | 5.36 | 0.157 | 68.79 | 3.13 |
| | 20.0 | F | 668 | 3.47 | 2.22 | 35.9 | 0.57 | 0.56 | 6.24 | 0.173 | 57.93 | 2.90 |
| | 19.0 | F | 710 | 3.02 | 1.70 | 43.8 | 0.60 | 0.59 | 5.10 | 0.159 | 49.76 | 2.62 |
| *Atf4*$^{cKO(e2)}$ | 20.0 | F | 682 | 3.66 | 2.35 | 35.8 | 0.61 | 0.63 | 5.82 | 0.183 | 71.83 | 3.59 |
| | 30.0 | M | 662 | 3.80 | 2.44 | 35.8 | 0.64 | 0.63 | 6.06 | 0.127 | 79.06 | 2.64 |
| | 29.0 | M | 634 | 3.66 | 2.32 | 36.6 | 0.65 | 0.65 | 5.64 | 0.126 | 76.66 | 2.64 |
| | 20.0 | F | 681 | 2.85 | 1.54 | 46.0 | 0.63 | 0.62 | 4.62 | 0.142 | 47.50 | 2.37 |

### Full unedited gels for:

Figure 2E

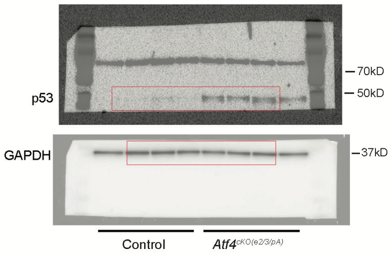


Figure 3D

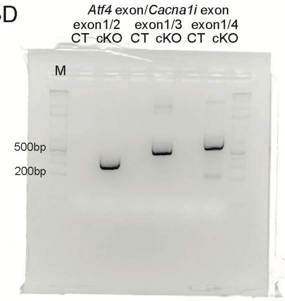


Figure 3E

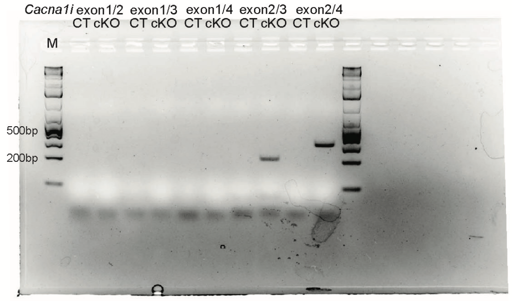


Figure 4B

Figure 4C

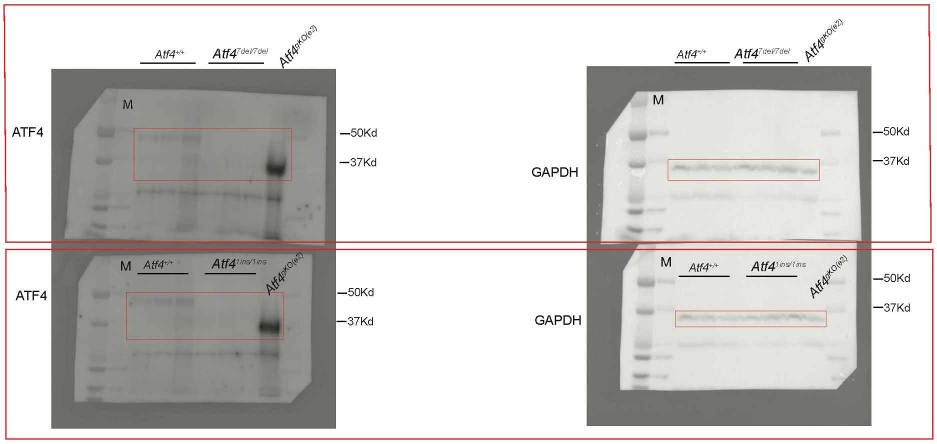


Figure 5F

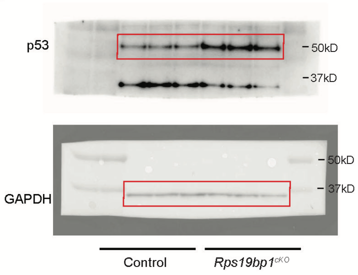

### Full unedited gels for:

Figure S4

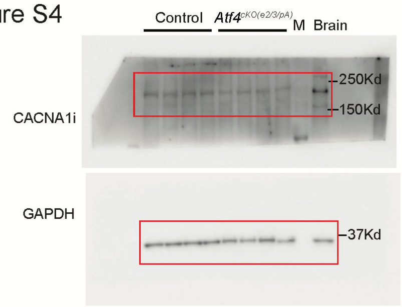


Figure S5

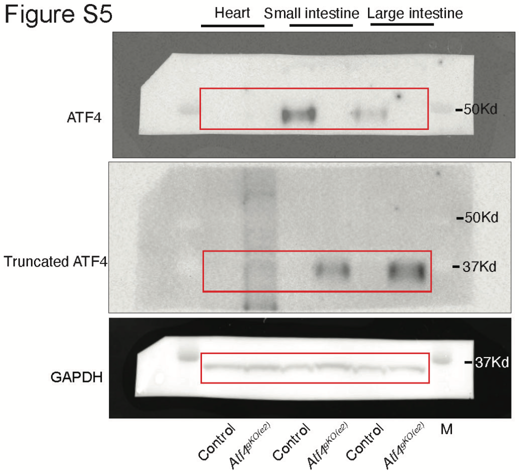
